# Pervasive off-target and double-stranded DNA nicking by CRISPR-Cas12a

**DOI:** 10.1101/657791

**Authors:** Karthik Murugan, Arun S. Seetharam, Andrew J. Severin, Dipali G. Sashital

## Abstract

Cas12a (formerly Cpf1) is an RNA-guided endonuclease in the CRISPR-Cas immune system that can be easily programmed for genome editing. Cas12a can bind and cut dsDNA targets with high specificity *in vivo*, making it an ideal candidate for precise genome editing applications. This specificity is contradictory to the natural role of Cas12a as an immune effector against rapidly evolving phages. However, the native cleavage specificity and activity remains to be fully understood. We employed high-throughput *in vitro* cleavage assays to determine and compare the native specificities of three Cas12a orthologs. Surprisingly, we observed pervasive nicking of randomized target libraries, with strong nicking activity observed against targets with up to four mismatches. Nicking and cleavage activities are dependent on mismatch type and position, and vary depending on the Cas12a ortholog and crRNA sequence. Our high-throughput and biochemical analysis further reveal that Cas12a has robust activated non-specific nicking and weak non-specific dsDNA degradation activity in trans. Together, our findings reveal Cas12a cleavage activities that could be beneficial in the context of bacterial CRISPR-Cas immunity but may be detrimental for genome editing technology.

## Introduction

Cas12a (formerly Cpf1) is an RNA-guided endonuclease that acts as the effector protein in type V-A CRISPR-Cas (clustered regularly interspaced short palindromic repeats-CRISPR associated) immune systems (Zetsche et al. 2015). Within these systems, CRISPR arrays are DNA loci consisting of unique sequences called spacers that are flanked by repeat sequences. Spacer sequences are acquired from foreign nucleic acids during infections and serve as memories to defend against future infections (Barrangou et al. 2007). The CRISPR locus is transcribed into a long pre-CRISPR RNA (pre-crRNA), which is processed by Cas12a into small mature CRISPR RNA (crRNA) (Fonfara et al. 2016). Cas12a uses these crRNAs as guides to bind to complementary “protospacer” sequences within the invading DNA (Zetsche et al. 2015). Following DNA binding, Cas12a sequentially cleaves each strand of the DNA using a RuvC nuclease domain, creating a double-strand break (DSB) that eventually leads to neutralization of the infection (Zetsche et al. 2015; Hille et al. 2018; Swarts and Jinek 2019). Because the crRNA can be changed to guide Cas12a to target a sequence of interest, Cas12a is easy to repurpose for programmable genome editing and other biotechnological applications (Zetsche et al. 2015), similar to the widely used *Streptococcus pyogenes* (Sp) Cas9 (Knott and Doudna 2018). Several orthologs, including *Francisella novicida* (Fn) Cas12a*, Lachnospiraceae bacterium* (Lb) Cas12a and *Acidaminococcus sp.* (As) Cas12a, have been used for genome editing (Zetsche et al. 2015, 2017).

The specificity of Cas endonucleases is an important consideration for both CRISPR-Cas immunity and genome editing applications. While SpCas9 has been shown to tolerate mutations in the target sequence (Hsu et al. 2013), Cas12a orthologs are reported to be relatively specific (D. Kim et al. 2016; Kleinstiver et al. 2016; H. K. Kim et al. 2017). In searching for potential targets, both Cas9 and Cas12a identify a protospacer adjacent motif (PAM) (Yamano et al. 2017; Sternberg et al. 2014) to initiate R-loop formation with the crRNA-complementary strand of the dsDNA target (Stella, Alcón, and Montoya 2017; Jiang et al. 2016). However, they differ in terms of structure and target cleavage mechanisms that may modulate target cleavage specificity (Swarts and Jinek 2018, 1; Murugan et al. 2017). The first few nucleotides following the PAM sequence (i.e. the PAM-proximal sequence) where DNA-RNA hybrid nucleation begins is referred to as the “seed” region (Swarts, Oost, and Jinek 2017). Mismatches in this region have been reported to be more deleterious for binding and cleavage by Cas12a than by SpCas9 (Hsu et al. 2013; Liu et al. 2016; Sternberg et al. 2014; Swarts, Oost, and Jinek 2017; Jeon et al. 2018). This intrinsic low tolerance for mismatches in the target sequence is a desirable trait for high-fidelity genome editing but raises the question of how Cas12a can provide effective immunity to bacteria against invaders in its native role. Higher mismatch tolerance may limit the ability of phages to escape from CRISPR-Cas immunity via mutations and may provide broader defense against closely related phages (Deveau et al. 2008; Tao, Wu, and Rao 2018). Recent reports showed that Cas12a can indiscriminately target single-stranded (ss) DNA in trans upon binding and activation by a crRNA-complementary dsDNA or ssDNA (Chen et al. 2018; Li et al. 2018). This activated, non-specific ssDNA cleavage may provide a wide-ranging immunity against ssDNA phage, even in the event of evolution. However, it remains unclear how the high specificity for dsDNA cleavage by Cas12a could defend against dsDNA phage escape via mutation.

In this study, we developed a high-throughput *in vitro* method to determine the native specificity and cleavage activity of Cas12a orthologs. We show that Cas12a nicks target sequences containing up to four mismatches, even though linearization does not always occur. We further show that Cas12a has robust activated non-specific nicking activity in trans, which could result in unpredicTable off-target nicking events during genome editing. Activated Cas12a also has weak dsDNA degradation activity for both target and non-specific DNA. Our results report several cleavage activities of Cas12a including cis (target-dependent) and activated trans (target-independent) dsDNA nicking, and cis and trans dsDNA degradation. These non-specific cleavage activities of Cas12a may aid bacteria containing type V-A CRISPR-Cas systems to fight against a variety of rapidly evolving phages.

## Results

### Cleavage activity of Cas12a against a target library

Genome editing activity and specificity of Cas12a have previously been characterized *in vivo* (D. Kim et al. 2016; Kleinstiver et al. 2016; H. K. Kim et al. 2017, 1). These studies show that Cas12a has low or no tolerance for mismatches in the target sequence in eukaryotic cells. However, the native cleavage specificity of Cas12a remains unclear, given that eukaryotic genomic structure may sequester potential off-targets (Hinz, Laughery, and Wyrick 2015; Isaac et al. 2016; Yarrington et al. 2018) and that most off-target analyses only account for the creation of double-strand breaks (Tsai et al. 2015, 2017). To directly observe the cleavage activity and specificity of Cas12a, we performed *in vitro* cleavage assays using a plasmid library (Fig. 1A). The libraries were designed to contain target sequences with 0 to 7 mismatches, with targets containing 2 or 3 mismatches maximally represented in the pool (“Pool Design, Complexity, and Purification” n.d.). The 0 mismatch “perfect” target was spiked in as an internal control (Supplementary Fig. 1A). We tested the cleavage activity of three Cas12a orthologs – FnCas12a, LbCas12a and AsCas12a (referred to collectively as Cas12a hereafter). For each Cas12a ortholog, we used two or three different crRNA sequences and corresponding negatively supercoiled plasmids containing the perfect target (pTarget) or target libraries (pLibrary) (Supplementary Fig. 1B – D, Supplementary Table 1). The three crRNA and library sequences were designed based on protospacer 4 sequence from *Streptococcus pyogenes* CRISPR locus (55% G/C), EMX1 gene target sequence (80% G/C) and CCR5 gene target sequence (20% G/C, tested for FnCas12a and AsCas12a), henceforth referred to as pLibrary PS4, EMX1 and CCR5 respectively.

**Figure 1:**
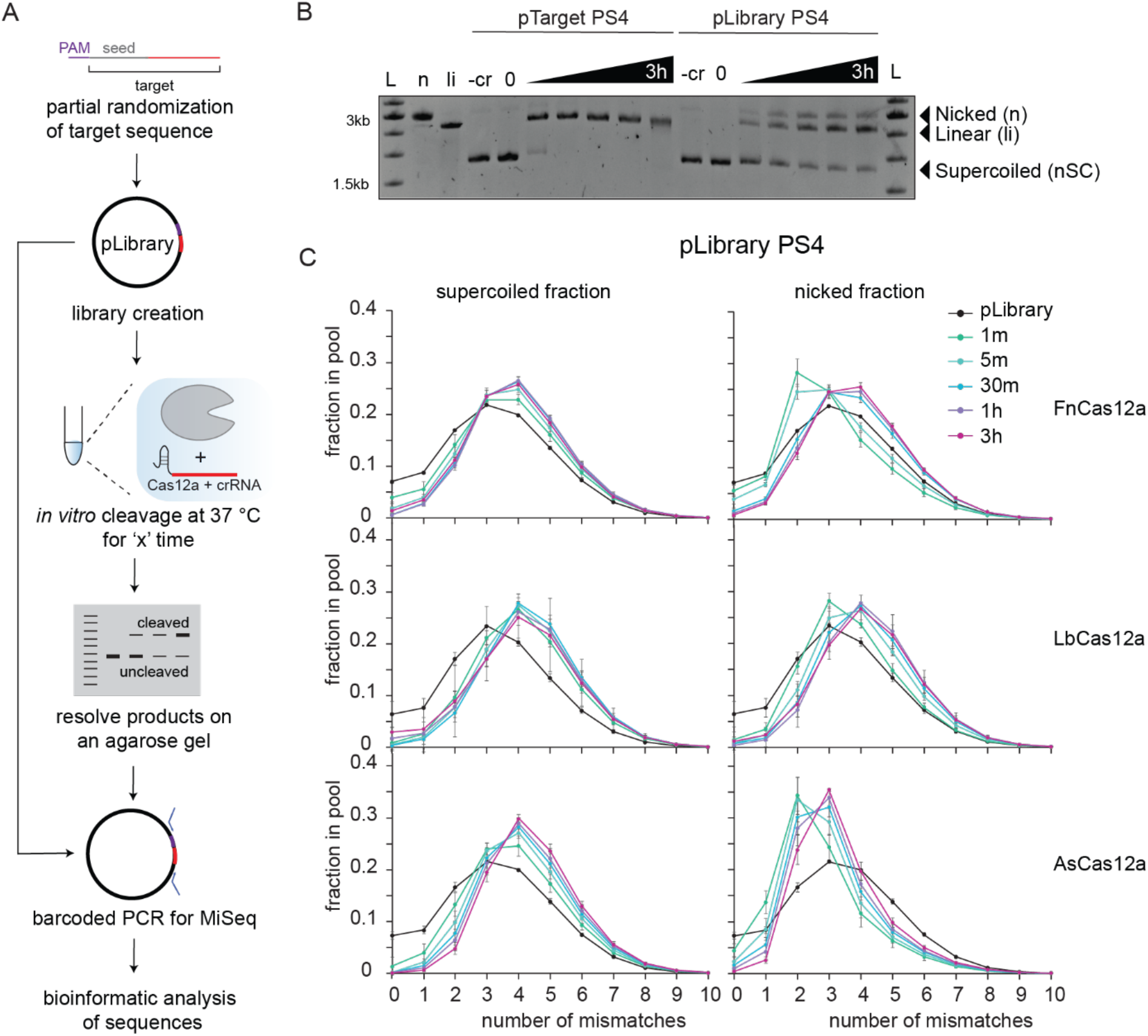
High-throughput *in vitro* cleavage assay reveals target-dependent cleavage and nicking activity of Cas12a. (A) Outline and workflow of the high-throughput *in vitro* cleavage assay. (B) Representative agarose gel showing time course cleavage of negatively supercoiled (nSC) plasmid containing a fully matched target (pTarget, left) and plasmid library (pLibrary, right) pLibrary PS4 by LbCas12a, resulting in linear (li) and/or nicked (n) products. Time points at which the samples were collected are 1 min, 5 min, 30 min, 1 hour, and 3 hours. Controls: -cr = reaction without cognate crRNA, n = Nt.BspQI nicked pUC19, li = BsaI-HF linearized pUC19 (C) Mismatch distribution of the pLibrary PS4 in the supercoiled and nicked fractions for different Cas12a orthologs. Depletion of target sequences from the supercoiled fraction indicates cleavage, and enrichment in the nicked fraction indicates nicking. The decrease in nicked fraction over time indicates linearization of target sequences. Error bars are SD, n = 3 replicates

In cleavage assays, Cas12a completely linearized the pTarget within the time course, indicating complete cleavage of both DNA strands (Fig. 1B, Supplementary Fig. 1E). In contrast, only a fraction of pLibrary was linearized and a substantial amount of plasmid remained supercoiled, indicating that many sequences within the pool could not be cleaved by Cas12a in the time span tested (Fig. 1B, Supplementary Fig. 1E). Surprisingly, we also observed a nicked fraction for pLibrary that persisted through the longest time point tested (3 hours) (Fig. 1B, Supplementary Fig. 1E).

To determine which sequences were uncleaved and nicked, we extracted the plasmid DNA from gel bands for each of these fractions, PCR amplified the target region and performed high-throughput sequencing (HTS) followed by bioinformatic analysis (Fig. 1A, Supplementary Fig. 2). Although we could not analyze sequences present in the linearized DNA sample, we assumed that sequences that were absent from both the supercoiled and nicked fractions were linearized. We generated mismatch distribution curves for the supercoiled and nicked fractions of the pLibraries subjected to Cas12a cleavage over time (Fig. 1C, Supplementary Fig. 1B – D, 3A, B). Depletion of target sequences from the supercoiled fraction indicates cleavage, and enrichment in the nicked fraction indicates nicking. The decrease in nicked fraction over time indicates linearization of target sequences. As expected, the perfect target (0 mismatch) was quickly depleted from the pool of sequences in the supercoiled and nicked fraction, and continued to decrease over time, indicating linearization (Fig. 1C, Supplementary Fig. 3A, B). For the target sequences with mismatches, the mismatch distribution of the pool shifted in comparison to the original pLibrary (Fig. 1C, Supplementary Fig. 1B – D, 3A, B).

Interestingly, Cas12a orthologs have differential cleavage activities against the three libraries tested. For pLibraries PS4 and EMX1, we observed rapid depletion of sequences with up to two mismatches from the supercoiled fractions within the first time point (1 min), indicating that all three Cas12a orthologs can tolerate up to 2 mismatches in these target sequences. We also consistently observed more depletion of sequences with three mismatches for LbCas12a and AsCas12a than for FnCas12a, suggesting that FnCas12a is less tolerant of these mismatches. Target sequences with 1 or 2 mismatches were depleted more rapidly for pLibrary CCR5 than the other two pLibraries, suggesting that mismatches can be tolerated better for A/T-rich sequences (Supplementary Fig. 3B). As with pLibraries PS4 and EMX1, sequences with three mismatches were depleted more rapidly from the pLibrary CCR5 supercoiled fraction for AsCas12a than for FnCas12a.

The mismatch distribution of the nicked fractions was not a uniform shift as observed for the supercoiled fractions (Fig 1C, Supplementary Fig. 3A, B). For most pLibraries subjected to FnCas12a and LbCas12a cleavage, target sequences with 3 or fewer mismatches were enriched in the nicked fraction in the first few time points but depleted over time, demonstrating that some of the mismatched sequences were eventually linearized. However, AsCas12a displayed less linearization activity against nicked targets, as target sequences with 2 or 3 mismatches were depleted more slowly or remained enriched in the nicked fraction over the entire time course for all pLibraries. We were also surprised to observe that FnCas12a and LbCas12a displayed strong nicking activity against target sequences containing 4 or more mismatches based on the enrichment of these sequences in the nicked fractions at the last time point tested (3 hours) for all pLibraries (Fig. 1C, Supplementary Fig. 3A, B). Overall, these results suggest that FnCas12a and LbCas12a are more tolerant of mismatches in the target sequences, and may be more prone to generating double-strand breaks at off-target sites than AsCas12a.

### Sequence determinants of Cas12a cleavage activity

We next looked at the sequences that were present in the supercoiled and nicked fractions to determine the effects of mismatch position and type (Supplementary Fig. 2). The heatmaps in Fig. 2 and Supplementary figures 4 – 10 show the relative distribution of target sequences containing 1 to 6 mismatches (MM) with all possible nucleotides at each position of the sequence in the supercoiled fraction, traced over time (see methods section – HTS analysis). Similar to the mismatch distribution of the supercoiled fraction for the pLibraries (Fig. 1C, Supplementary Fig. 3A, B), target sequences with a single mismatch (1 MM) were quickly depleted from the supercoiled fraction over time for all pLibraries tested (Fig. 2, Supplementary Fig. 4 – 10). Some mismatches in the PAM-proximal “seed” region were enriched in the supercoiled fraction and/or depleted slowly, indicating that these mismatches were not tolerated by Cas12a. We observed a short seed region of ~6 nucleotides for most pLibraries and Cas12a orthologs. This is in agreement with previously reported *in vivo* and *in vitro* specificity of Cas12a (D. Kim et al. 2016; Kleinstiver et al. 2016; H. K. Kim et al. 2017; Swarts, Oost, and Jinek 2017), and is shorter compared the ~10 nucleotide length for SpCas9 (Hsu et al. 2013; Liu et al. 2016; Sternberg et al. 2014). A single G substitution in the seed region is highly deleterious for cleavage by Cas12a, while C or T mismatches slowed the rate of cleavage or depletion from the supercoiled fractions to a lesser degree (Fig 2, Supplementary Fig. 4 – 10). In contrast, target sequences with a single A mismatch in the seed are generally tolerated for cleavage outside of the first PAM-proximal position for most pLibraries. Outside of the seed, most target sequences containing a single mismatch were rapidly depleted from the supercoiled fraction, indicating that any type of single mismatch outside the seed can be similarly tolerated. However, in some cases single mismatches throughout the target sequence slow the rate of cleavage, most notably for EMX1 pLibrary cleavage by LbCas12a and FnCas12a (Supplementary Fig. 6 – 8).

**Figure 2:**
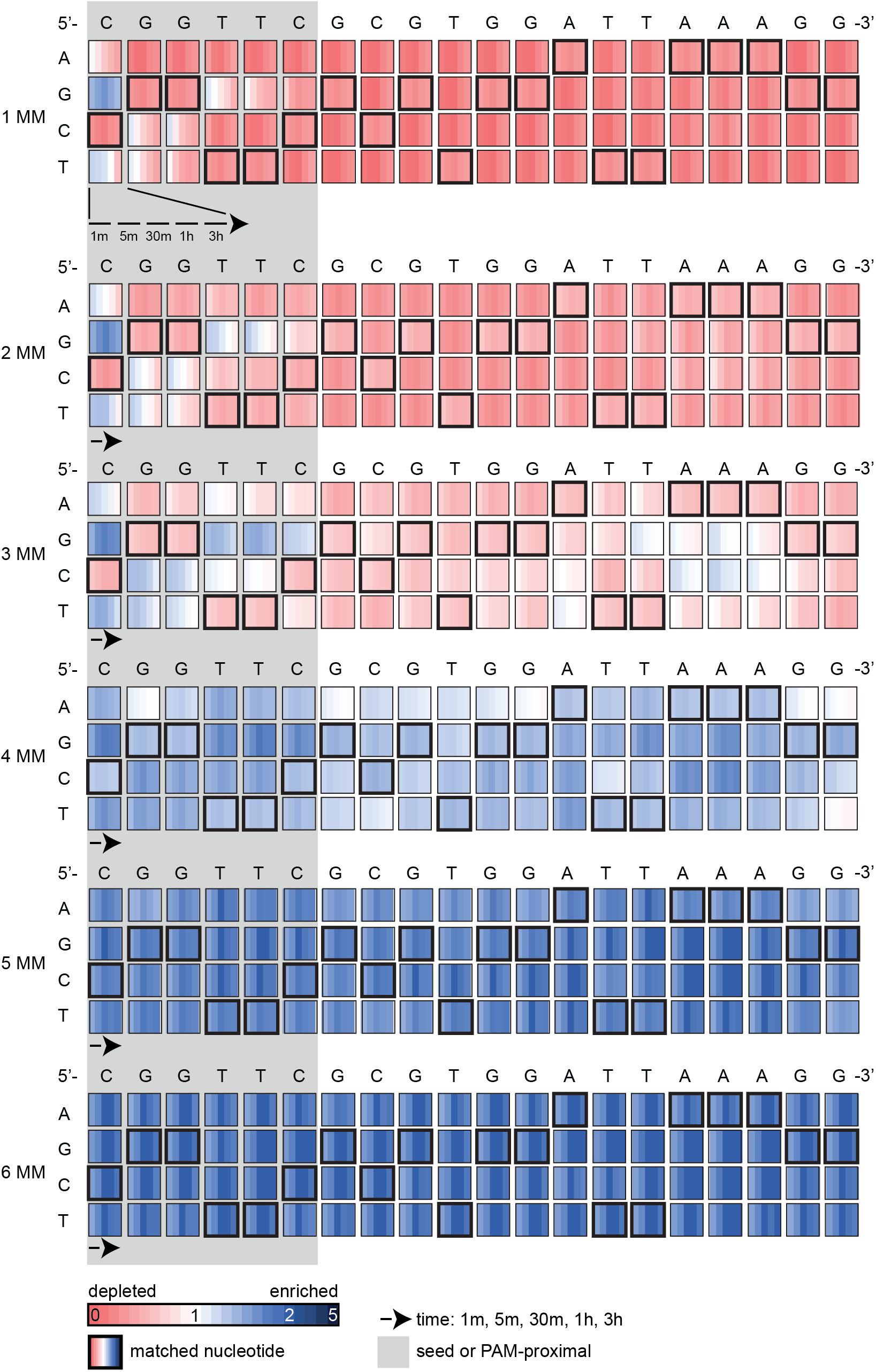
Sequence determinants of LbCas12a cleavage activity for pLibrary PS4. Heatmaps showing the relative enrichment (blue) or depletion (red) of different mismatched sequences over time for the supercoiled fraction in pLibrary PS4 upon cleavage by LbCas12a. The target sequence is indicated on the top and highlighted by bold black boxes in the heatmap. The PAM-proximal “seed” sequence is highlighted by the grey box. Each box represents the proportion of sequences containing each nucleotide at a given position across time. Values plotted represent average of three replicates. MM = mismatch

A similar seed-dependent cleavage trend was observed for target sequences with 2, 3 and 4 mismatches (2 MM, 3 MM and 4 MM, respectively) (Fig 2, Supplementary Fig. 4 – 10). In general, G substitution is most deleterious for cleavage at all positions in the target sequence with 3 and 4 mismatches. T and C substitutions also slow the rate of cleavage by Cas12a, particularly in pLibrary CCR5 (Supplementary Fig. 9, 10). For LbCas12a, most target sequences with 2 and 3 mismatches in the PAM-distal regions were eventually depleted in pLibraries PS4 and EMX1 (Fig. 2, Supplementary Fig. 7). FnCas12a tolerates most transition and transversion substitutions in the PAM-distal region of target sequences with 2 and 3 mismatches across all three pLibraries (Supplementary Fig. 4, 6, 9). AsCas12a tolerates up to 3 mismatches in the PAM-distal region for pLibrary CCR5 (Supplementary Fig. 10), as seen in the library mismatch distribution of the supercoiled fraction (Supplementary Fig. 3B). For sequences with 5 and 6 mismatches, we observed a steady enrichment of all sequences regardless of mismatch position or type, indicating that Cas12a does not have sequence-specific cleavage activity against these targets (Fig 2, Supplementary Fig. 4 – 10).

### Sequence determinants of Cas12a nickase activity

To determine sequences that were preferentially nicked by Cas12a, we performed analysis on the nicked fraction similar to the supercoiled fraction. The heatmaps in Fig. 3 and Supplementary figures 11 – 17 show the relative distribution of target sequences containing 1 to 6 mismatches (MM) with all possible nucleotides at each position of the sequence in the nicked fraction, traced over time (see methods section – HTS analysis).

**Figure 3:**
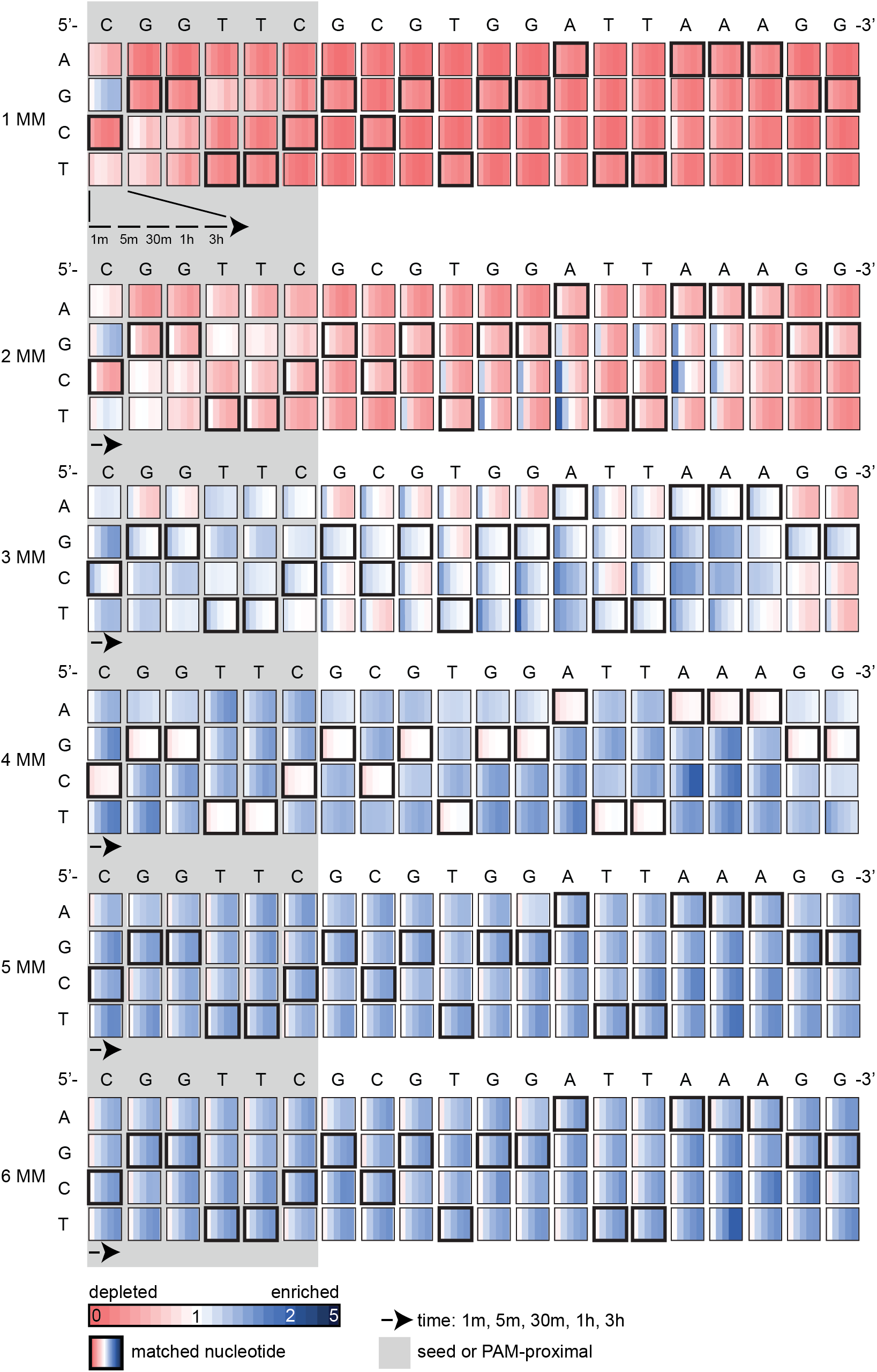
LbCas12a has pervasive nicking activity against mismatched sequences in pLibrary PS4. Heatmaps showing the relative enrichment (blue) or depletion (red) of different mismatched sequences over time for the nicked fraction in pLibrary PS4 upon cleavage by LbCas12a. The target sequence is indicated on the top and highlighted by bold black boxes in the heatmap. The PAM-proximal “seed” sequence is highlighted by the grey box. Each box represents the proportion of sequences containing each nucleotide at a given position across time. Values plotted represent average of three replicates. MM = mismatch

Cas12a linearized most single mismatches (1 MM) within the time frame tested (3 hours) (Fig. 3, Supplementary Fig 11 – 17). While LbCas12a rapidly linearized target sequences with most single mismatches, FnCas12a and AsCas12a displayed slower kinetics for double-stranded cleavage for some single mismatch sequences. Interestingly, several of these deleterious mismatches were located in the region where the non-target strand is cut (position 16 - 18 from the 5’ end of the target on the non-target strand) (Zetsche et al. 2015; Strohkendl et al. 2018) (Supplementary Fig. 11, 12, 15, 17). Similarly, target sequences with 2 and 3 mismatches (2 MM and 3 MM, respectively) in the PAM-distal region were enriched in the nicked fraction at early time points (Fig. 3, Supplementary Fig 11 – 17). For LbCas12a and FnCas12a, these 2 and 3 mismatch targets were eventually depleted, indicating that these mismatches are tolerated for linearization (Fig. 3, Supplementary Fig. 11, 13, 14, 16). For AsCas12a cleavage of PS4 and EMX1 pLibraries, target sequences with 2 and 3 mismatches in the PAM-distal region remained highly enriched throughout the time course (Supplementary Fig. 12, 15, 17), indicating that mismatches in the PAM-distal region block the second cleavage step for this Cas12a ortholog. Taken together, these data suggest that LbCas12a and FnCas12a can linearize most target sequences with 2 and 3 mismatches while AsCas12a can only nick these target sequences. These results support our observations that FnCas12a and LbCas12a are more tolerant to mismatches than AsCas12a for double-stranded cleavage.

FnCas12a and LbCas12a can nick target sequences with 4 or more mismatches, observed as a strong enrichment of these target sequences for almost all pLibraries tested (Fig. 3, Supplementary Fig 11 – 17). Similar to the supercoiled fraction, target sequences with 5 and 6 mismatches were uniformly enriched in the nicked fraction irrespective of the mismatch position or type for most pLibraries and Cas12a orthologs (Fig 3, Supplementary Fig. 11 – 17). However, AsCas12a did not nick target sequences with 4 or more mismatches for pLibraries PS4 and EMX1 (Supplementary Fig. 12, 15), but exhibited weak nicking of these target sequences in pLibrary CCR5 (Supplementary Fig. 17).

### Cas12a has non-specific nicking and dsDNA degradation activity

Our HTS data suggests that Cas12a can nick sequences with several mismatches. To validate this observation, we selected sequences from pLibrary PS4 that were relatively enriched in the nicked fraction at the longest time point (3 hours). We cloned sequences containing 2 to 8 mismatches and individually tested Cas12a nicking activity against each plasmid. Consistent with our HTS results, we observed varying degrees of nicking and linearization of target sequences containing 2, 3 and 4 mismatches (MM) for different Cas12a orthologs (Fig. 4A). To compare among the Cas12a orthologs, we quantified the supercoiled, linearized and nicked fractions at the 3-hour time point (based on the time frame used for the pLibrary cleavage) for the perfect and mismatched target sequences (Fig. 4B, Supplementary Fig. 18B, C). FnCas12a and AsCas12a showed higher target-dependent nicking activity, while LbCas12a partially linearized most targets. All three Cas12a orthologs strongly nicked one of the 3 mismatch target sequences (3.2 MM) tested, with no or low partial linearization of the mismatched target.

**Figure 4:**
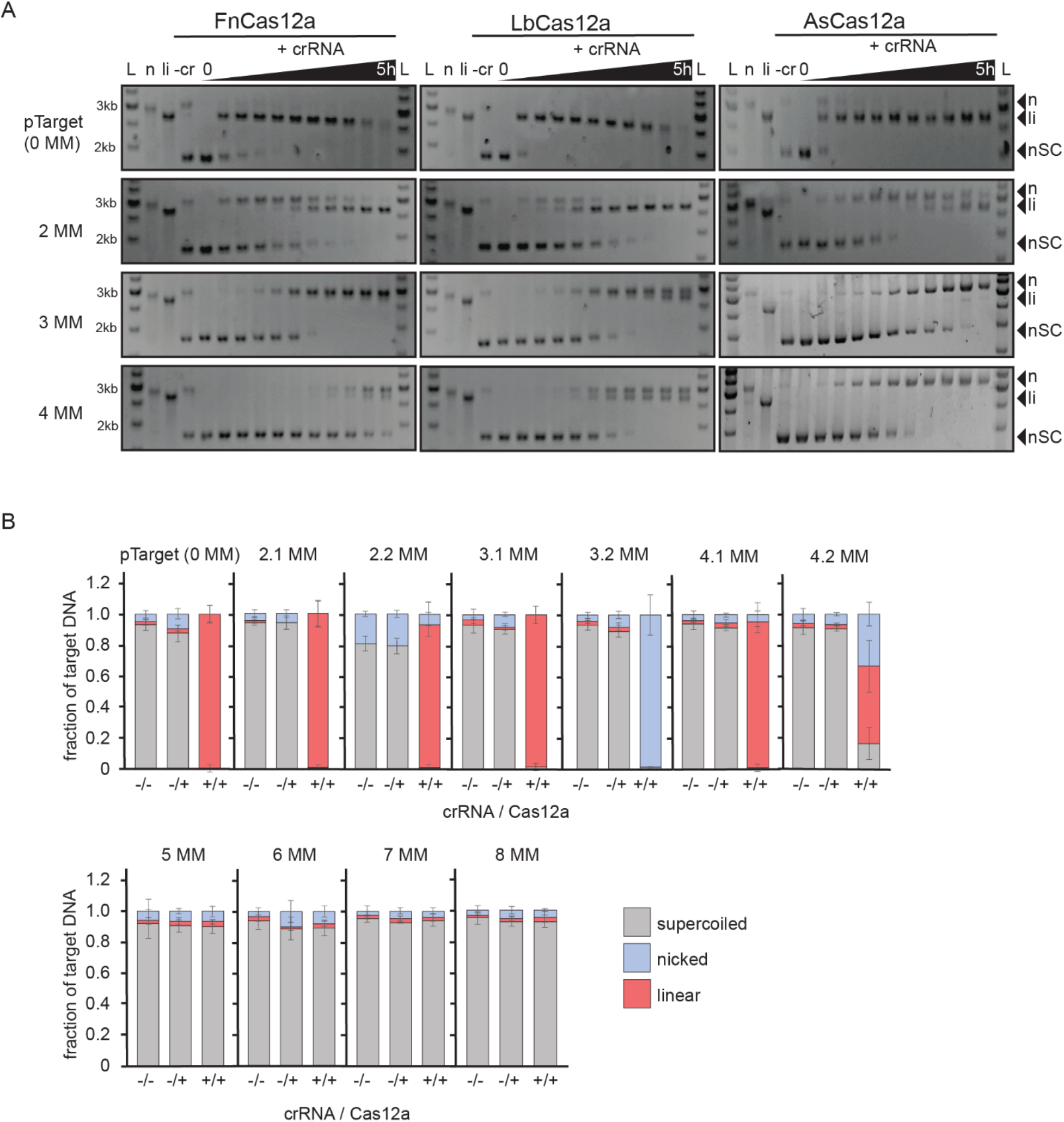
Cas12a orthologs have distinct nicking patterns against mismatched targets. (A) Representative agarose gels showing cleavage of a negatively supercoiled (nSC) plasmid containing the perfect target (pTarget) or mismatched (MM) target over a time course by Cas12a orthologs, FnCas12a (left), LbCas12a (center) and AsCas12a (right), resulting in linear (li) and/or nicked (n) products. Time points at which the samples were collected are 15 sec, 30 sec, 1 min, 2 min, 5 min, 15 min, 30 min, 1 hour, 3 hours, and 5 hours. Controls: -cr = reaction without cognate crRNA, n = Nt.BspQI nicked pUC19, li = BsaI-HF linearized pUC19. (B) Quantification of supercoiled, linear and nicked fractions from cleavage of perfect and mismatched (MM) target plasmid by LbCas12a after 3 hours. -/- indicates a cleavage reaction with the target plasmid without Cas12a and crRNA. -/+ indicates a cleavage reaction with the target plasmid and Cas12a only, and +/+ indicates a cleavage reaction with the target plasmid, Cas12a and cognate crRNA. Average of the intensity fraction values are plotted with SD as error bars, n = 3 replicates.

Surprisingly, sequences with greater than five mismatches were not nicked by Cas12a, although these sequences were enriched in the nicked fraction of our HTS data (Fig. 4B, Supplementary Fig. 18A – C). Notably, heatmaps for target sequences with 5 and 6 mismatches indicated enrichment of these sequences in the nicked fraction was sequence non-specific (Fig. 3, Supplementary Fig. 11, 13, 14, 16, 17). This led us to hypothesize that Cas12a may have target-activated non-specific nicking activity against targets with low or no homology to the crRNA. In the pLibrary cleavage assays, the mixed pool of sequences contains the perfect target sequence which may activate Cas12a for non-specific nicking activity (Chen et al. 2018; Li et al. 2018). To test this, we used a short dsDNA oligonucleotide activator that was fully complementary to the crRNA to activate Cas12a. We formed a complex containing Cas12a, crRNA and the dsDNA activator and tested for cleavage activity against empty plasmid in three forms – negatively supercoiled (nSC), nicked (n) and linear (li). Surprisingly, we observed robust non-specific, trans nicking and partial linearization of the empty negatively supercoiled plasmid by Cas12a (Fig. 5A, Supplementary Fig. 19A). To confirm that the non-specific nicking activity was not due to an artifact in our protein purification, we also tested commercially available LbCas12a (New England Biolabs, NEB LbCas12a) and AsCas12a (Integrated DNA Technologies, IDT AsCas12a). We observed that the non-specific, activator-mediated nicking activity was reproducible with these commercial enzymes (Supplementary Fig. 19A). This activity was also reproducible with other crRNA-activator pairs, although the activation of the trans nicking activity varied with the crRNA-activator pair (Supplementary Fig. 20). Taken together, these results demonstrate that Cas12a can be activated for non-specific nicking of dsDNA targets.

**Figure 5:**
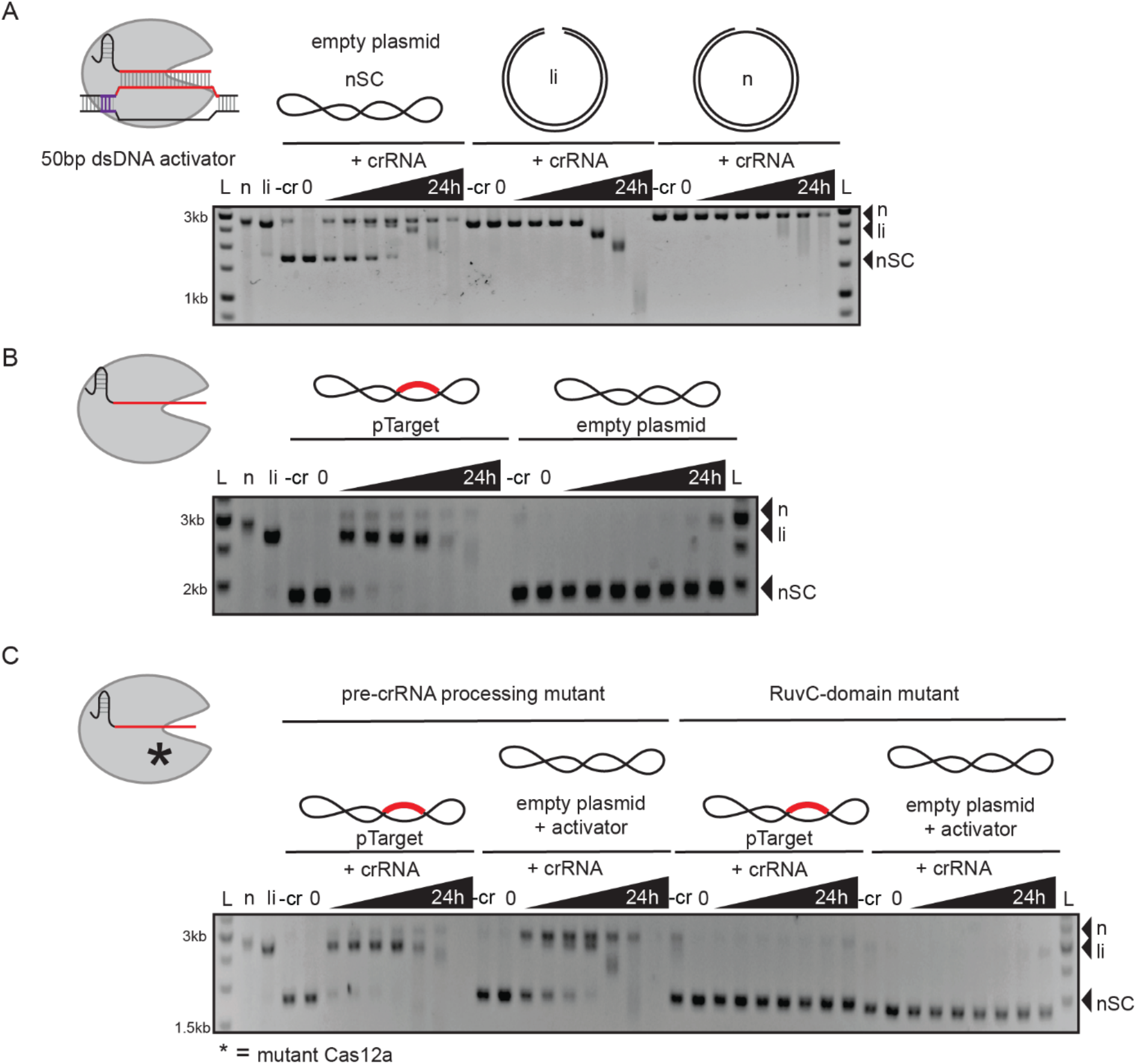
Cas12a has activated non-specific, trans dsDNA nicking and degradation activity. (A) Representative agarose gel showing non-specific nicking and linearization of negatively supercoiled (nSC) dsDNA plasmid, and degradation of linearized and nicked dsDNA plasmid over time by LbCas12a. LbCas12a (20 nM) and crRNA (30 nM) were complexed with a crRNA-complementary 50 bp dsDNA activator (30 nM) (perfect target from pLibrary PS4) and incubated with non-specific dsDNA in different forms – negatively supercoiled (nSC), linear (li) and nicked (n). (B) Representative agarose gel showing target-dependent cleavage and degradation of dsDNA by LbCas12a. (C) Representative agarose gel showing RuvC-domain dependent pTarget linearization and activated non-specific nicking and degradation of dsDNA plasmid. Pre-crRNA processing and RuvC-domain active site mutants of LbCas12a (20 nM) and crRNA (30 nM) were complexed with a crRNA-complementary 50bp dsDNA activator (30 nM) (perfect target from pLibrary PS4) and incubated with non-specific dsDNA in different forms – negatively supercoiled (nSC), linear (li) and nicked (n). Time points at which the samples were collected are 5 min, 15 min, 30 min, 1 hour, 4 hours, 8 hours, and 24 hours. Controls: -cr = reaction without cognate crRNA, n = Nt.BspQI nicked pUC19, li = BsaI-HF linearized pUC19.

While both FnCas12a and LbCas12a have strong non-specific nicking activity, AsCas12a is not strongly activated as a nickase upon target binding (Supplementary Fig. 19A). The lack of AsCas12a activated nickase activity is evident in the HTS data, where sequences containing more than 4 mismatches were not strongly enriched in the nicked fraction (Supplementary Fig. 12, 15, 17). The reduced activation could be due to slower rates of PAM-distal product release after cleavage, where the cleaved products hinder RuvC-domain from accessing other dsDNA substrates (Swarts and Jinek 2019; Singh et al. 2018).

In addition to non-specific nicking by activated Cas12a, we also observed linearization of the nicked dsDNA over time and degradation of empty linear and nicked plasmid (Fig. 5A, Supplementary Fig. 19A). To further investigate this activity, we first tested cleavage of pTarget by Cas12a and performed the cleavage assay for longer time points (up to 24h). We observed slow, processive degradation of the DNA target plasmid after 4 hours of incubation with Cas12a (Fig. 5B, Supplementary Fig. 19B). Similarly, target-activated Cas12a can degrade non-specific dsDNA after nicking and linearizing it (Fig. 5A, Supplementary Fig. 19A). Like the non-specific nicking and ssDNase activity (Chen et al. 2018; Li et al. 2018), Cas12a can be activated by crRNA-complementary ssDNA binding in a PAM-independent, RuvC-domain dependent manner for non-specific dsDNA nicking and degradation (Fig. 5C, Supplementary Fig. 19C, 21A, B).

We next tested whether mismatched target sequences that were present in the pLibrary cleavage assays could also act as activators. Interestingly, Cas12a was activated by some of these mismatched targets as well, especially those that were partially linearized by Cas12a (Supplementary Fig. 22A). However, mismatched targets that were only nicked by Cas12a (Fig. 4B, Supplementary Fig. 18B, C) were weak activators, indicating that double-strand cleavage of the target activator is important for activated dsDNA nicking, as has been previously observed for target-activated ssDNA degradation (Chen et al. 2018; Li et al. 2018; Swarts and Jinek 2019).

## Discussion

Cas12a has become a widely used tool for various biotechnological applications such as genome editing and diagnostic tools (Zetsche et al. 2017; Chen et al. 2018; Gootenberg et al. 2018). Several reports show that Cas12a and engineered orthologs are highly specific for RNA-guided dsDNA cleavage activity (D. Kim et al. 2016; Kleinstiver et al. 2016; H. K. Kim et al. 2017). Despite these studies on Cas12a specificity, the cleavage activity and specificity outside of a eukaryotic setting remains unclear. The apparent high specificity of Cas12a in genome editing studies is paradoxical to its natural role as an immune system effector. Phages evolve rapidly and can escape from CRISPR-Cas immunity via mutations (Deveau et al. 2008; Tao, Wu, and Rao 2018). The high specificity of Cas12a may also limit targeting of closely related phages (Andersson and Banfield 2008).

Here we show that Cas12a has additional dsDNA nicking and degradation activities apart from previously described crRNA-mediated cis cleavage of dsDNA targets (Zetsche et al. 2015, 1) and activated trans cleavage of non-specific ssDNA substrates (Chen et al. 2018; Li et al. 2018). Our results demonstrate that Cas12a can nick and, in some cases, create double-strand breaks in targets with up to four mismatches. Similarly, a recent study by Fu et al. (Fu et al. 2019) demonstrated that Cas12a and Cas9 have target-dependent nicking activity against targets with one or two mismatches. We also establish the Cas12a has non-specific dsDNA nicking activity upon binding to a crRNA-complementary DNA. While this manuscript was in preparation, a complementary study reported similar observations, demonstrating that these activities are reproducible *in vitro* (Fig. 6) (Fuchs et al. 2019). It is also interesting to note that several proteins in the Cas12 family have strong nickase activities against target DNA, but can only weakly linearize dsDNA targets (Yan et al. 2019; Strecker et al. 2019).

**Figure 6:**
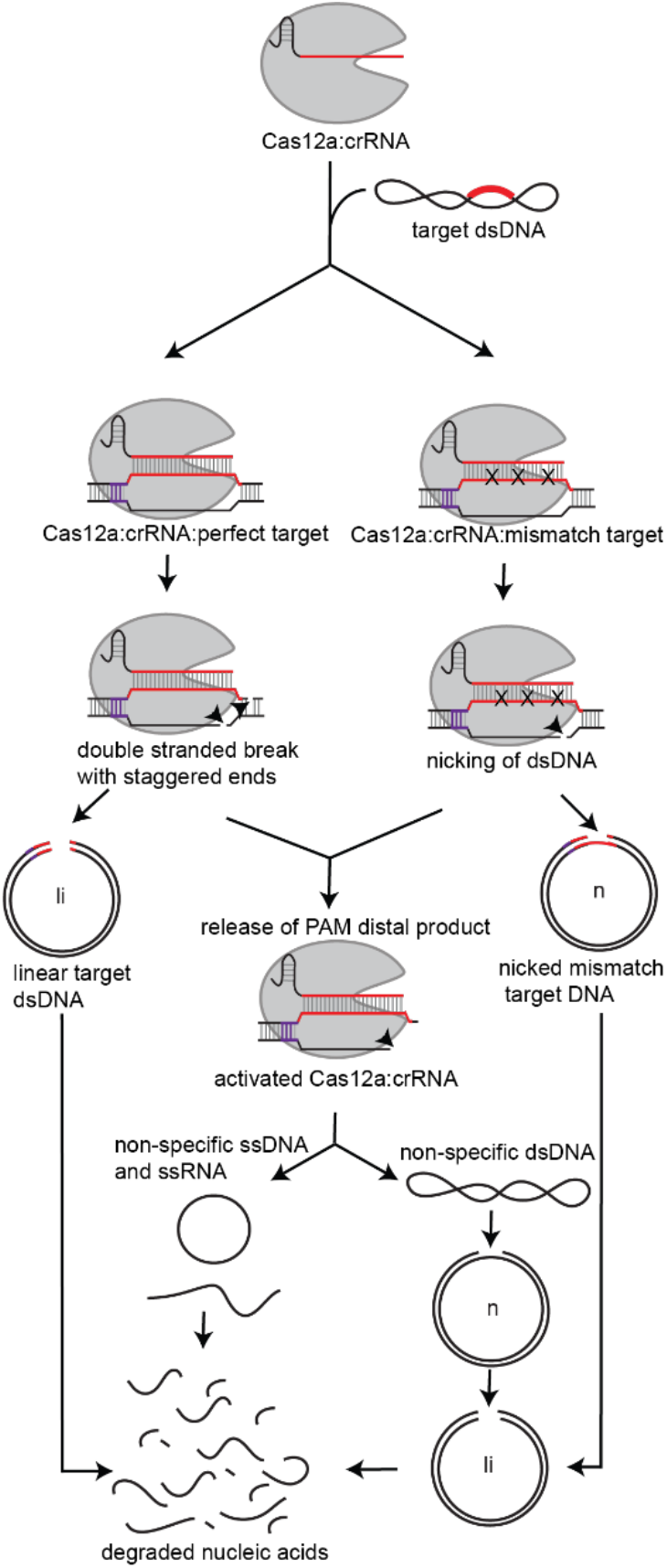
Cas12a has cis and activated, trans nuclease activities. Cas12a-crRNA complex can bind complementary target dsDNA and cleave it. Some mismatched targets can only be nicked by Cas12a. After this cleavage event, Cas12a releases the PAM distal cleavage product and remains bound to at least the PAM proximal cleaved target-strand, thus remaining in an activated confirmation. The active RuvC domain can then accept other nucleic acid substrates. Target-activated Cas12a can further nick, linearize and degrade non-specific dsDNA, ssDNA and ssRNA substrates.

Cas12a has a single active site, and it cleaves the dsDNA target in a sequential order with the nicking of non-target strand (NTS) followed by target-strand (TS) (Swarts and Jinek 2019; Jeon et al. 2018). As a result, Cas12a cleaves the NTS faster than the TS (Strohkendl et al. 2018; Stella et al. 2018). Single mismatches slow the rate of R-loop formation by Cas12a, but it remains unknown how multiple mismatches in the target sequence affect the cleavage mechanism. The difference in rates of strand cleavage could potentially be amplified by the presence of mismatches in the target sequence, resulting in a nicked intermediate. By testing three different pLibraries, we show the effects of different nucleotide substitutions across the target region in varying mismatch combinations and that some mismatches are not tolerated for cleavage while others can be nicked. Some mismatches slow the linearization of the target sequence, suggesting that specific types of mismatches may cause slower cleavage of the TS. In this case, TS cleavage may be the rate limiting step. Another possibility is that the target dissociates following NTS cleavage due to R-loop collapse or that the TS is not presented to the RuvC domain. Biochemical studies have revealed that the TS recognition is a pre-requisite for NTS and ssDNA cleavage (Chen et al. 2018). However, NTS cleavage can occur even when the TS is modified and uncleavable but not the other way around (Swarts and Jinek 2019). The release of the PAM-distal products upon cleavage exposes the active RuvC domain that can accept substrates for non-specific trans cleavage activities (Singh et al. 2018; Swarts and Jinek 2019; Jeon et al. 2018). Interestingly, we observed that certain mismatched targets can only be nicked by Cas12a and these natural mismatched substrates are ineffective activators of the non-specific nicking activity of Cas12a presumably due to failed cleavage and release of PAM-distal TS products.

A recent study demonstrated that topology plays a role in Cas12a-mediated target DNA cleavage (van Aelst et al. 2019). Similarly, we observed different rates of nicking and degradation of different forms of DNA. Nicking by activated-Cas12a was easily detected for negatively supercoiled DNA. Cas12a is likely to mainly encounter negatively supercoiled DNA in both prokaryotic and eukaryotic cells, suggesting that the activated trans cleavage could enable dsDNA nicking *in vivo*. Cas12a can also slowly degrade linear dsDNA. This can be attributed to the fraying of DNA at the exposed sites (Andreatta et al. 2006; Fei and Ha 2013). The exposed ssDNA regions are likely degraded via the ssDNase activity of Cas12a (Chen et al. 2018; Li et al. 2018). Multiple events of the trans nicking activity may also cause the DNA to fall apart (Fuchs et al. 2019). This may be true in the case of nicked dsDNA substrates where we observed direct degradation rather than an intermediate linear product. This suggests that a cumulative effect of all the trans activities of Cas12a eventually leads to complete degradation of nucleic acid substrates (Fig. 6).

Cas12a has been successfully used for gene editing *in vivo* without any deleterious off-target effects (D. Kim et al. 2016; Kleinstiver et al. 2016; H. K. Kim et al. 2017; Tang et al. 2018; Moon et al. 2018). Our high-throughput pLibrary cleavage analysis indicates that Cas12a can bind and cleave sequences with up to four mismatches *in vitro*. However, the cellular context also plays a role in Cas effector binding and cleavage. SpCas9 can bind to targets containing several mismatches depending on DNA breathing and supercoiling, both *in vitro* and *in vivo* (Newton et al. 2019; Wu et al. 2014). Cas12a can stably bind to targets with mismatches *in vitro* (Singh et al. 2018), but *in vivo* studies suggest low or no off-target binding (Zhang et al. 2017). This could reflect the inability of Cas12a to unwind and bind DNA in varying topological and cellular contexts which may result in overall lower off-target editing rates by Cas12a.

While off-target sites can be predicted and avoided by careful design of crRNAs, the robust non-specific nicking activity we observed *in vitro* may lead to off-target editing as nicked DNA can recruit DNA repair machinery *in vivo* (Kuzminov 2001; Vriend et al. 2016; Vriend and Krawczyk 2017), although nicks may be repaired by error-free DNA repair pathways (Fukui 2010). Nevertheless, the unpredicTable nature of the non-specific nicking activity makes it difficult to detect the outcomes of nicking. In addition, the commonly used methods to verify off-target editing do not detect nicks (Tsai et al. 2015, 2017), meaning that detection of potential off-target effects due to non-specific nicking could require whole genome sequencing. Use of Cas12a orthologs, such as AsCas12a, that display reduced non-specific nickase activity may reduce these unpredictable effects during genome editing experiments. Notably, our *in vitro* specificity analysis also suggests that AsCas12a is less prone to creating double-strand breaks at sites with two or three mismatches, suggesting that this ortholog may be less prone to off-target cleavage at highly homologous sites.

The activities reported in our study add to the growing number of specific and non-specific Cas12a cleavage activities (Fig. 6) and may provide a possible explanation of how Cas12a compensates for its highly specific targeted cleavage activities as an immune effector. The target-dependent non-specific nicking and degradation activities, along with previously described dsDNA cleavage (Zetsche et al. 2015) and trans ssDNase activity (Chen et al. 2018; Li et al. 2018), could allow Cas12a to mount a strong defense against different types of invading phages. In the event of phage evolution via mutations, Cas12a may tolerate some mutations and still nick or fully cleave phage DNA. Cas12a could also be activated by the evolved target region of the phage DNA, enabling non-specific nicking and degradation activities. The non-specific nature of the nicking and degradation activities may also be harmful to the host bacteria. In CRISPR-Cas systems like type III and type VI, Cas nucleases can be activated for non-specific cleavage of RNA (Hille et al. 2018). Perhaps, this is a means to prevent phage proliferation by initiating programmed cell death (PCD), abortive infection or dormancy in order to save the bacterial population (Koonin and Zhang 2017; Meeske, Nakandakari-Higa, and Marraffini 2019). Further studies are required to investigate the cost-benefit relation of such non-specific activities of Cas12a to the bacteria.

## Acknowledgements

We thank all the former and current members of the Sashital Lab for helpful discussions and suggestions on various aspects of the project. We thank Michael Baker from the DNA Facility for assistance with HTS data collection, and the Protein Facility for providing access to ImageQuant TL, both a branch of the Iowa State University Office of Biotechnology. We also thank Heather S. Lewin and Megan N. O’Donnell from the University Library for helping with the data deposition to DataShare.

This work was supported by funds from startup funds to D.G.S from Iowa State University College of Liberal Arts and Sciences and the Roy J. Carver ChariTable Trust, the National Science Foundation (1652661 to D.G.S), and National Institute of Food and Agriculture (IOW05480 to D.G.S).

## Author contributions

All experiments and HTS sample preparation were performed by K.M. The HTS data extraction was performed by A.S.S and A.J.S. HTS results were analyzed and interpreted by K.M and D.G.S. The manuscript was written by K.M and D.G.S, with input from A.S.S and A.J.S. All authors read the final draft of the manuscript. Funding for this project was secured by D.G.S.

## Data Availability

HTS data and processed data files from this study have been deposited in the Iowa State University Library’s DataShare, and can be found at https://doi.org/10.25380/iastate.8178938. All other information and data are available from the authors upon request.

## Methods

### Cas12a cloning

The gene sequences for *Francisella novicida* (Fn) and *Acidaminococcus sp* (As) Cas12a were synthesized as *Escherichia coli* codon-optimized gBlocks (purchased from Integrated DNA Technologies, IDT) and inserted into pSV272 using Gibson assembly (New England Biolabs) as per the manufacturer’s protocol, to generate a protein expression construct encoding Cas12a fused with N-terminal 6X-His sequence, a maltose binding protein (MBP) and a Tobacco Etch Virus (TEV) protease cleavage site*. Lachnospiraceae bacterium* (Lb) Cas12a was expressed using expression plasmid pMAL-his-LbCpf1-EC. pMAL-his-LbCpf1-EC was a gift from Jin-Soo Kim (Addgene plasmid # 79008; http://n2t.net/addgene:79008; RRID:Addgene_79008) (D. Kim et al. 2016). Catalytically inactive (pre-crRNA processing and DNase dead) Cas12a mutants were generated via site-directed mutagenesis (SDM) and verified by Sanger sequencing (Eurofins Genomics, Kentucky, USA) (see Supplementary Table 1 for SDM primers).

### Cas12a expression and purification

Purification protocols were adapted from previously established Cas12a purification methods (Mohanraju et al. 2018). All Cas12a proteins were expressed in *Escherichia coli* BL21 (DE3) cells. 2X TY broth supplemented with corresponding antibiotics was inoculated with overnight cultures of cells in 1:100 ratio. Cultures were grown to an optical density (600 nm) of 0.5 − 0.6 at 37 °C and protein expression was induced by the addition of IPTG to a final concentration of 0.2 mM. The incubation was continued at 18 °C overnight.

For LbCas12a, AsCas12a and all Cas12a mutants, cells were harvested by centrifugation and the cell pellet was resuspended in Lysis Buffer I (20 mM Tris-HCl pH 8.0, 500 mM NaCl, 5 mM imidazole), supplemented with protease inhibitors PMSF, cOmplete™ Protease Inhibitor Cocktail Tablet or Halt Protease Inhibitor Cocktail. Cells were lysed by sonication or a homogenizer and the lysate was centrifuged to remove insoluble material. The clarified lysate was applied to a HisPur™ Ni-NTA Resin (ThermoFisher Scientific) column. The column was washed with 10 column volumes of Wash Buffer (Lysis Buffer + 15 mM imidazole final concentration) and bound protein was eluted in Elution Buffer I (20 mM Tris-HCl pH 8.0, 500 mM NaCl, 250 mM imidazole). Fractions containing Cas12a were pooled and TEV protease was added in a 1:100 (w/w) ratio. The sample was dialyzed in Dialysis Buffer (10 mM HEPES-KOH pH 7.5, 200 mM KCl, 1 mM DTT) at 4°C overnight. For further purification the protein was diluted 1:1 with 20 mM HEPES KOH (pH 7.5) and loaded on a HiTrap Heparin HP (GE Healthcare) column. The column was washed with Buffer A (20 mM HEPES-KOH pH 7.5, 100 mM KCl) and eluted with Buffer B (20 mM HEPES-KOH pH 7.5, 2 M KCl) by applying a gradient from 0% to 50% over a total volume of 60 ml. Peak fractions were analyzed by SDS-PAGE and fractions containing Cas12a were combined, and DTT was added to a final concentration of 1 mM. The protein was fractionated on a HiLoad 16/600 Superdex 200 gel filtration column (GE Healthcare), eluting with SEC buffer (20 mM HEPES-KOH pH 7.5, 500 mM KCl, 1mM DTT). Peak fractions were combined, concentrated, flash frozen in liquid nitrogen and stored at −80°C until further use.

FnCas12a was purified by the following protocol. Cells were resuspended in Lysis Buffer II (20 mM Tris-HCl pH 8.0, 500 mM NaCl, 10mM imidazole, and 10% glycerol) supplemented with PMSF. Cells were lysed by sonication or a homogenizer and the lysate was centrifuged to remove insoluble material. The clarified lysate was applied to a HisPur™ Ni-NTA Resin (ThermoFisher Scientific) column. The column was washed with 10 column volumes of Lysis Buffer and bound protein was eluted in Elution Buffer II (Lysis Buffer II + 250 mM imidazole final concentration). The elution concentrated and run on a HiLoad 16/600 Superdex 200 gel filtration column (GE Healthcare) pre-equilibrated with SEC Buffer A (20 mM Tris-HCl, pH 8.0, and 500 mM NaCl). Fractions containing 6X His-MBP tagged Cas12a were collected and treated with TEV protease in a 1:100 (w/w) ratio, overnight at 4 °C. Samples were reapplied to HisPur™ Ni-NTA Resin (ThermoFisher Scientific) to remove the His-tagged TEV, free 6X His-MBP, and any remaining tagged protein. The flow-through was collected, concentrated and further purified by using a HiLoad 16/600 S200 gel filtration column in SEC Buffer B (20 mM Tris-HCl, pH 8.0, 200 mM KCl, and 1mM EDTA). Peak fractions were combined, concentrated, flash frozen in liquid nitrogen and stored at −80°C until further use.

Commercially available Cas12a were purchased from New England Biolabs (NEB) – LbCas12a and Integrated DNA Technologies (IDT) – AsCas12a.

### Library creation

To generate a pool of sequences containing mismatches, the library was partially randomized (“Pool Design, Complexity, and Purification” n.d.). The following probability distribution function was used to determine the randomization/doping frequency,

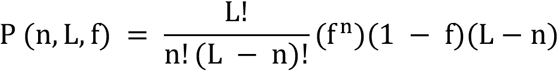

where, P is the fraction of the population, L is the sequence length, n is the number of mutations/template and f is the probability of mutation/position (doping level or frequency). The three library sequences tested were a modified protospacer 4 sequence from *Streptococcus pyogenes* CRISPR locus (55% GC), EMX1 gene target sequence (80% GC) and CCR5 gene target sequence (20% GC) (see Supplementary Table 1 for target sequence). A randomization/doping frequency (f) of 15% was selected to optimize the library to contain a maximum of sequences with 2 or 3 mismatches.

### Library preparation

Single-stranded oligonucleotides with 15% doping frequency in the target region were ordered from Integrated DNA Technologies (IDT). The oligonucleotides were diluted to 0.2 µM in 1X NEBuffer 2. pUC19 vector was amplified via PCR to insert homology arms. 30 ng of PCR amplified pUC19, 5 µL of library oligonucleotide (0.2 µM) and ddH_2_O to bring the volume 10 µL were mixed with 10 µL 2X NEBuilder HiFi DNA Assembly Master mix (New England Biolabs) and incubated at 50 °C for 1 h. 2 µL of the assembled product was transformed into NEB STable competent cells as per the manufacturer’s protocol. After the recovery step, all of cells in the outgrowth media were used to inoculate a 50 mL LB supplemented with ampicillin and incubated overnight at 37 °C. Cells were harvested and the plasmid library (pLibrary) was extracted using QIAGEN Plasmid Midi Kit.

### DNA and RNA preparation

DNA oligonucleotides were synthesized by IDT or Thermo Scientific. All RNAs were ordered from IDT. Sequences of DNA oligonucleotides and RNA used are in Supplementary Table 1. Target and mismatched target oligonucleotides were inserted into pUC19 using Gibson Assembly as described in the library preparation method. Target plasmids and empty pUC19 were linearized by restriction enzyme digestion using BsaI-HF and nicked using a nicking enzyme Nt.BspQI (New England Biolabs). All restriction digestion and Gibson Assembly reactions were carried out as per the manufacturer’s protocols. All sequences were verified by Sanger sequencing (Eurofins Genomics, Kentucky, USA).

### Plasmid DNA Cleavage Assay

The protocol was adapted from previously described methods (Anders and Jinek 2014). Briefly, Cas12a:crRNA complex was formed by incubating Cas12a and crRNA (1:1.5 ratio) in 1X CutSmart buffer (50 mM Potassium Acetate, 20 mM Tris-acetate, 10 mM Magnesium Acetate, 100 μg/ml BSA, pH 7.9) and 5 mM DTT at 37 °C for 10 min. For activator-mediated cleavage assays, Cas12a, crRNA, and dsDNA oligonucleotide activator (1:1.5:1.5) were incubated at 37 °C for 10 min. Cleavage reactions were initiated by mixing the Cas12a complex with pTarget or pLibrary (150 ng) and incubating at 37 °C. 10 µL aliquots were drawn from the reaction at each time point and quenched with phenol-chloroform. The aqueous layer was extracted and separated on a 1% agarose gel via electrophoresis and stained with SYBR safe or RED safe stain for visualization. Bands were visualized and quantified with ImageQuant TL (GE Heathcare). Intensities of the band (I) in the uncleaved (supercoiled - SC) and cleaved fractions (nicked - N and linearized - L) were measured. Fraction cleaved and uncleaved were calculated as follows.

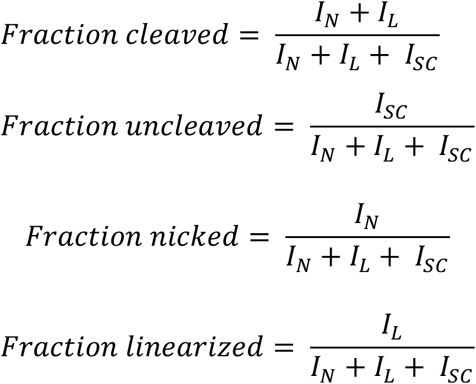

For library and mismatched target plasmid cleavage assays, 100 nM Cas12a and 150 nM crRNA was used. Concentrations of pLibrary, pTarget and empty pUC19 used were at 150 ng/10 µL (~7 nM) reaction. For activator-mediated cleavage assays, 20 nM Cas12a, 30 nM crRNA and 30 nM activator DNA oligonucleotide were used.

### Library preparation for HTS

The library plasmid cleavage products were run on an agarose gel as described above to separate the cleaved (linear and nicked) and uncleaved (supercoiled) products. The bands from the nicked and supercoiled fractions from various time points were excised and gel purified using QIAquick Gel Extraction Kit (Qiagen). PCR was used to add Nextera Adapters, followed by another round to add unique indices/barcodes for each sample. Sample size and quality were verified using DNA 1000 kit and Agilent 2100 Bioanalyzer. Samples were sent for MiSeq or NextSeq for paired-end reads of 150 cycles to Iowa State DNA Facility or Admera Health, LLC (New Jersey, USA). 15% PhiX was spiked in.

### HTS analysis

HTS data were obtained as compressed fastq files and were processed with custom bash scripts (see associated GitHub repository https://github.com/sashital-lab/Cas12a_nickase). A simple workflow of the analysis is described in Supplementary Fig. 2. Briefly, the files were renamed based on the sample information (pLibrary name, replicate and Cas12a ortholog), stored in separate folders identified by the library, and the target sequences were extracted. Bash scripts were used for obtaining the counts of the extracted target sequences, determining the number of mismatches, calculating the fractions in each replicate, as well as preparing summary Tables for total counts of each mismatched target sequence. Once all the processing was done on the command-line, they were imported in to Microsoft Excel for plotting and summarizing.

The fraction of target sequences containing ‘n’ mismatches (MM) (F_n-MM_) in the pool was calculated as:

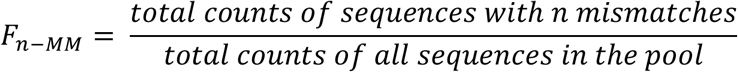

The relative change (enrichment and/or depletion) (R_c_) of a sequence ‘S’ containing ‘n’ mismatches at each time point ‘x’ compared to the control pLibrary was calculated as:

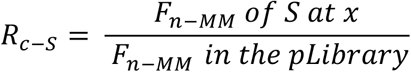

