## Supplemental material for "Pervasive off-target and double-stranded DNA nicking by CRISPR-Cas12a"

**This file includes:**

Supplementary Figures S1 to S22

Supplementary Table 1: List of Oligonucleotides

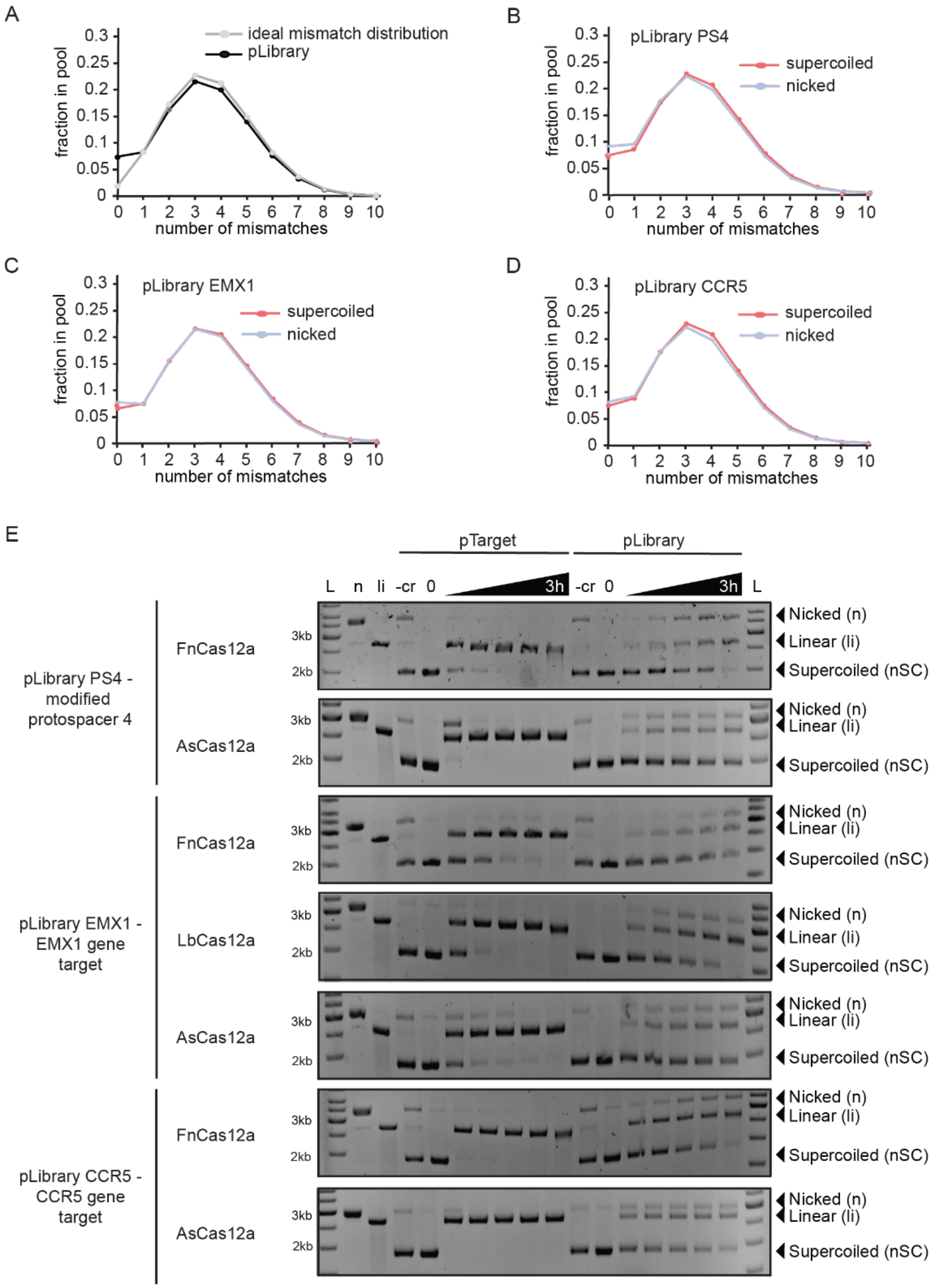


**Supplementary Figure 1: Target cleavage and nicking by Cas12a observed for different plasmid libraries.**

(A) Mismatch distribution of ideal library (gray) and experimental pLibrary (black) with a randomization or doping frequency (f) of 15%. The ideal mismatch distribution was calculated based on the formula shown in Methods section (Library creation). The doping frequency was selected to create a library that contained a maximum representation of sequences with 2 or 3 mismatches. The 0 mismatch “perfect” target was spiked into pLibrary as an internal control, resulting in a slight change from the ideal distribution.

(B – C) Mismatch distribution of the supercoiled and nicked fractions from different plasmid library controls, (B) pLibrary PS4, (C) pLibrary EMX1, (D) pLibrary CCR5. Although a clear nicked plasmid band was not visible in the gel with SYBR or RED safe staining, a band excised from the gel in the region where the nicked fraction would run produced a similar mismatch distribution to the supercoiled fraction when subjected to HTS, indicating the presence of trace amounts of nicked pLibrary prior to Cas12a cleavage.

(E) Agarose gel showing time course cleavage of negatively supercoiled (nSC) plasmid containing a target (pTarget, left) and plasmid library (pLibrary, right) by different Cas12a orthologs, resulting in linear (li) and/or nicked (n) products. Three different library sequences were tested for Cas12a cleavage activity, pLibrary PS4 (modified protospacer 4), EMX1 and CCR5 gene target (see methods – library creation and Supplementary Table 1).

Time points at which the samples were collected are 1 min, 5 min, 30 min, 1 hour, and 3 hours.

Controls: -cr = reaction without cognate crRNA, n = Nt.BspQI nicked pUC19, li = BsaI-HF linearized pUC19

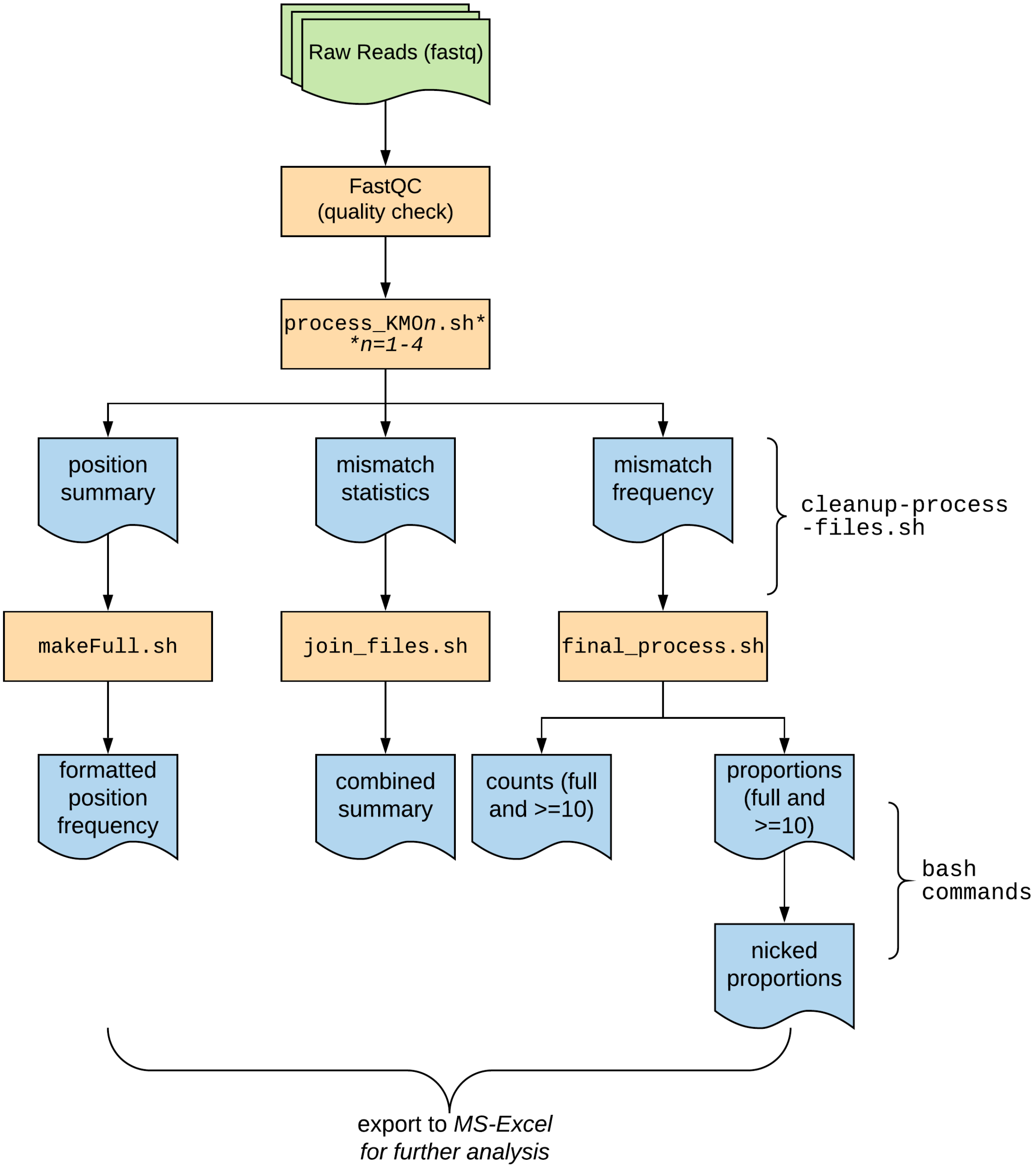


**Supplementary Figure 2: Workflow of the bioinformatic analysis of the HTS data.**
Target sequences were extracted from the HTS data using custom scripts. The mismatch number and position were determined and Tables reporting the counts and fractions of each mismatched target sequence were generated for further analysis (see methods – HTS data analysis).

A

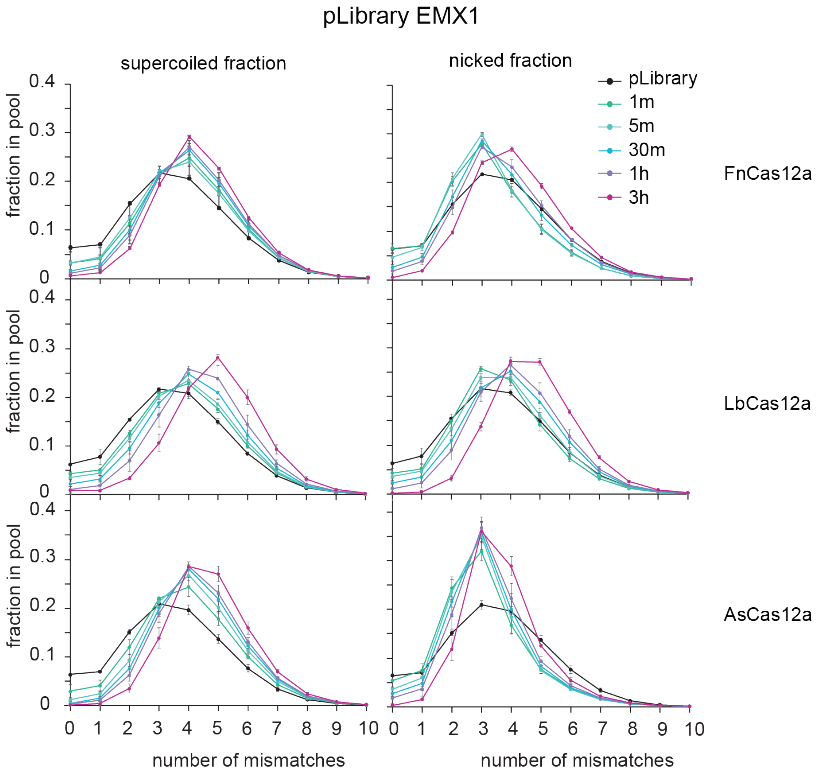


B

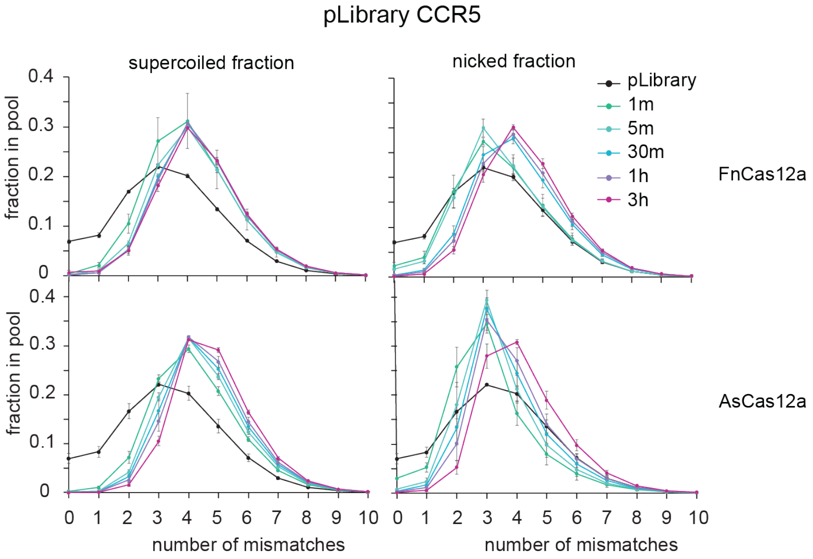


**Supplementary Figure 3: Target-dependent cleavage and nicking activity of Cas12a on two pLibraries.**

Mismatch distribution of two pLibraries (A) EMX1 and (B) CCR5 when subject to cleavage by different Cas12a orthologs. Depletion of target sequences from the supercoiled fraction indicates cleavage, and enrichment in the nicked fraction indicates nicking. The decrease in nicked fraction over time indicates linearization of target sequences. Error bars are SD, n = 3 replicates

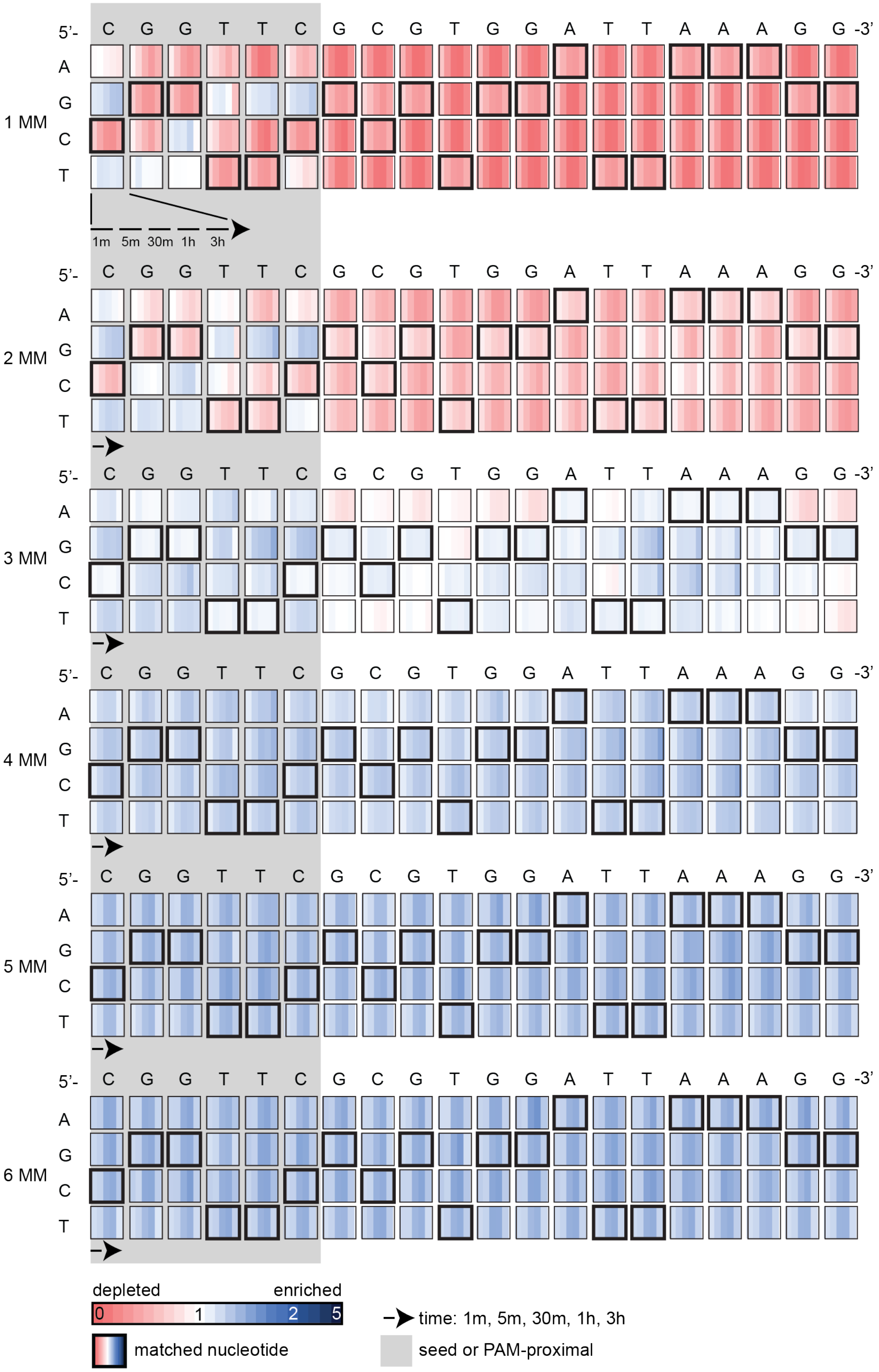

5'- C G G T T C G C G T G G A T T A A A G G -3'
A
G
C
T
1 MM
2 MM
3 MM
4 MM
5 MM
6 MM
1m 5m 30m 1h 3h
depleted
enriched
0 1 2 5
matched nucleotide
time: 1m, 5m, 30m, 1h, 3h
seed or PAM-proximal

**Supplementary Figure 4: Mismatched target sequences in the supercoiled fraction of pLibrary PS4 upon cleavage by FnCas12a.**

Heatmaps showing the relative enrichment (blue) or depletion (red) of different mismatched sequences over time for the supercoiled fraction in pLibrary PS4 upon cleavage by FnCas12a. The target sequence is indicated on the top and highlighted by bold black boxes in the heatmap. The PAM-proximal “seed” sequence is highlighted by the grey box. Each box represents the proportion of sequences containing each nucleotide at a given position across time. Values plotted represent average of three replicates. MM = mismatch

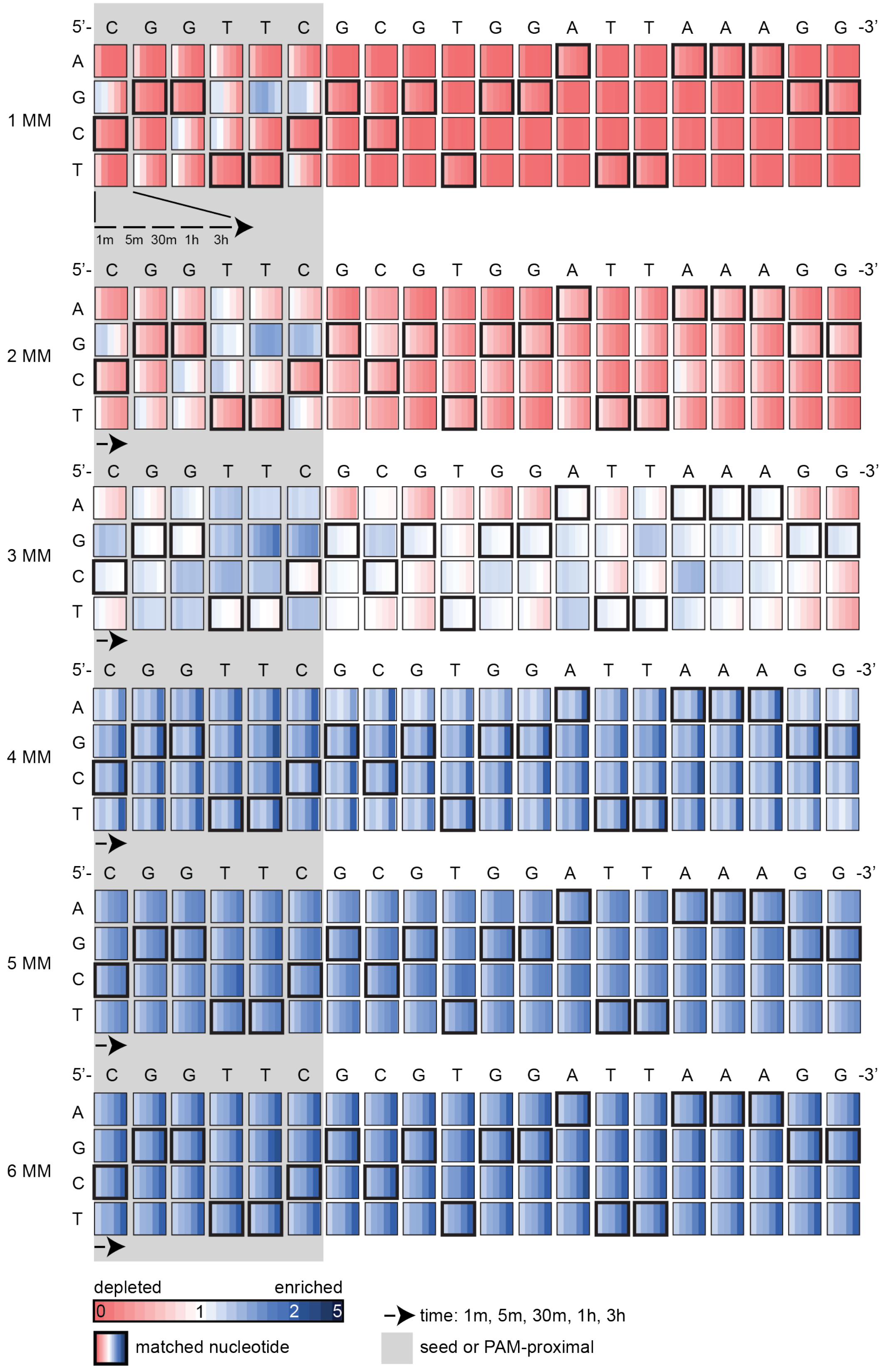

5'- C G G T T C G C G T G G A T T A A A G G -3'
A
G
C
T
1 MM
2 MM
3 MM
4 MM
5 MM
6 MM
1m 5m 30m 1h 3h
depleted
enriched
0
1
2
5
matched nucleotide
time: 1m, 5m, 30m, 1h, 3h
seed or PAM-proximal

**Supplementary Figure 5: Mismatched target sequences in the supercoiled fraction of pLibrary PS4 upon cleavage by AsCas12a.**

Heatmaps showing the relative enrichment (blue) or depletion (red) of different mismatched sequences over time for the supercoiled fraction in pLibrary PS4 upon cleavage by AsCas12a. The target sequence is indicated on the top and highlighted by bold black boxes in the heatmap. The PAM-proximal "seed" sequence is highlighted by the grey box. Each box represents the proportion of sequences containing each nucleotide at a given position across time. Values plotted represent average of three replicates. MM = mismatch

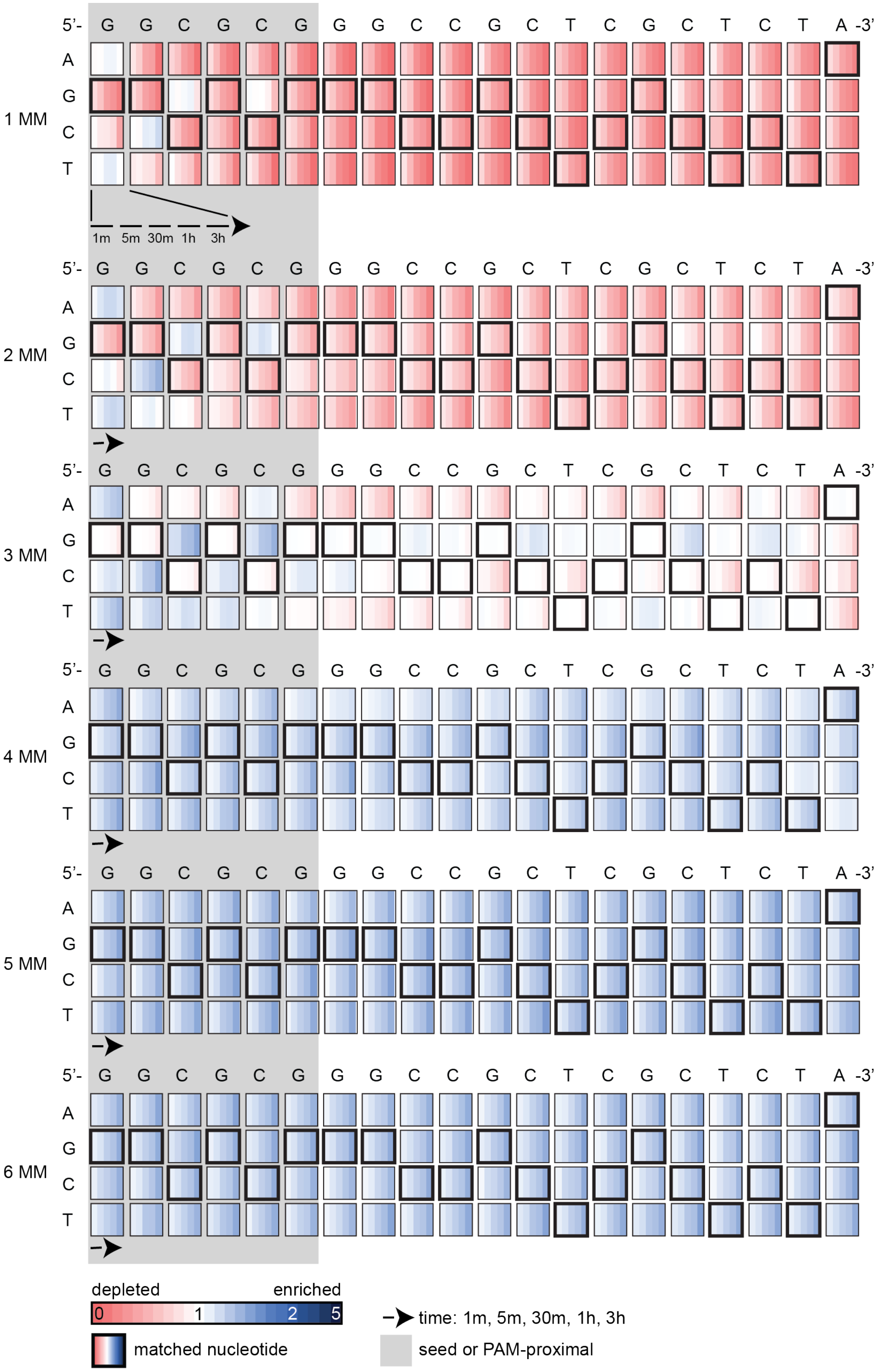
5'- G G C G C G G G C C G C T C G C T C T A -3'
A
G
C
T
1 MM
2 MM
3 MM
4 MM
5 MM
6 MM
1m 5m 30m 1h 3h
depleted
enriched
0
1
2
5
matched nucleotide
time: 1m, 5m, 30m, 1h, 3h
seed or PAM-proximal

**Supplementary Figure 6: Mismatched target sequences in the supercoiled fraction of pLibrary EMX1 upon cleavage by FnCas12a.**

Heatmaps showing the relative enrichment (blue) or depletion (red) of different mismatched sequences over time for the supercoiled fraction in pLibrary EMX1 upon cleavage by FnCas12a. The target sequence is indicated on the top and highlighted by bold black boxes in the heatmap. The PAM-proximal “seed” sequence is highlighted by the grey box. Each box represents the proportion of sequences containing each nucleotide at a given position across time. Values plotted represent average of three replicates. MM = mismatch

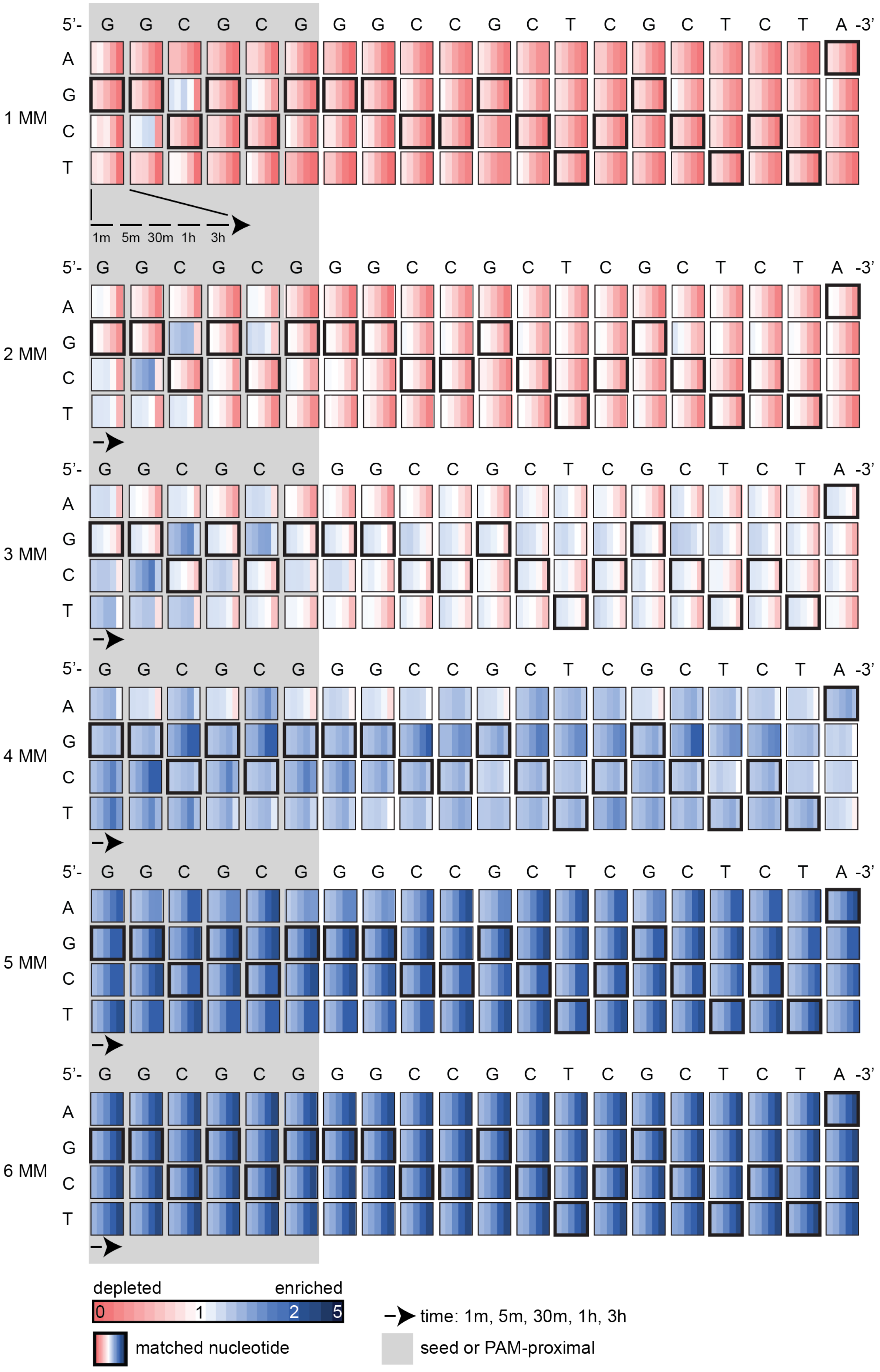
5'- G G C G C G G G C C G C T C G C T C T A -3'
A
G
C
T
1 MM
1m 5m 30m 1h 3h
2 MM
3 MM
4 MM
5 MM
6 MM
depleted
enriched
0
1
2
5
matched nucleotide
time: 1m, 5m, 30m, 1h, 3h
seed or PAM-proximal

**Supplementary Figure 7: Mismatched target sequences in the supercoiled fraction of pLibrary EMX1 upon cleavage by LbCas12a.**

Heatmaps showing the relative enrichment (blue) or depletion (red) of different mismatched sequences over time for the supercoiled fraction in pLibrary EMX1 upon cleavage by LbCas12a. The target sequence is indicated on the top and highlighted by bold black boxes in the heatmap. The PAM-proximal "seed" sequence is highlighted by the grey box. Each box represents the proportion of sequences containing each nucleotide at a given position across time. Values plotted represent average of three replicates. MM = mismatch

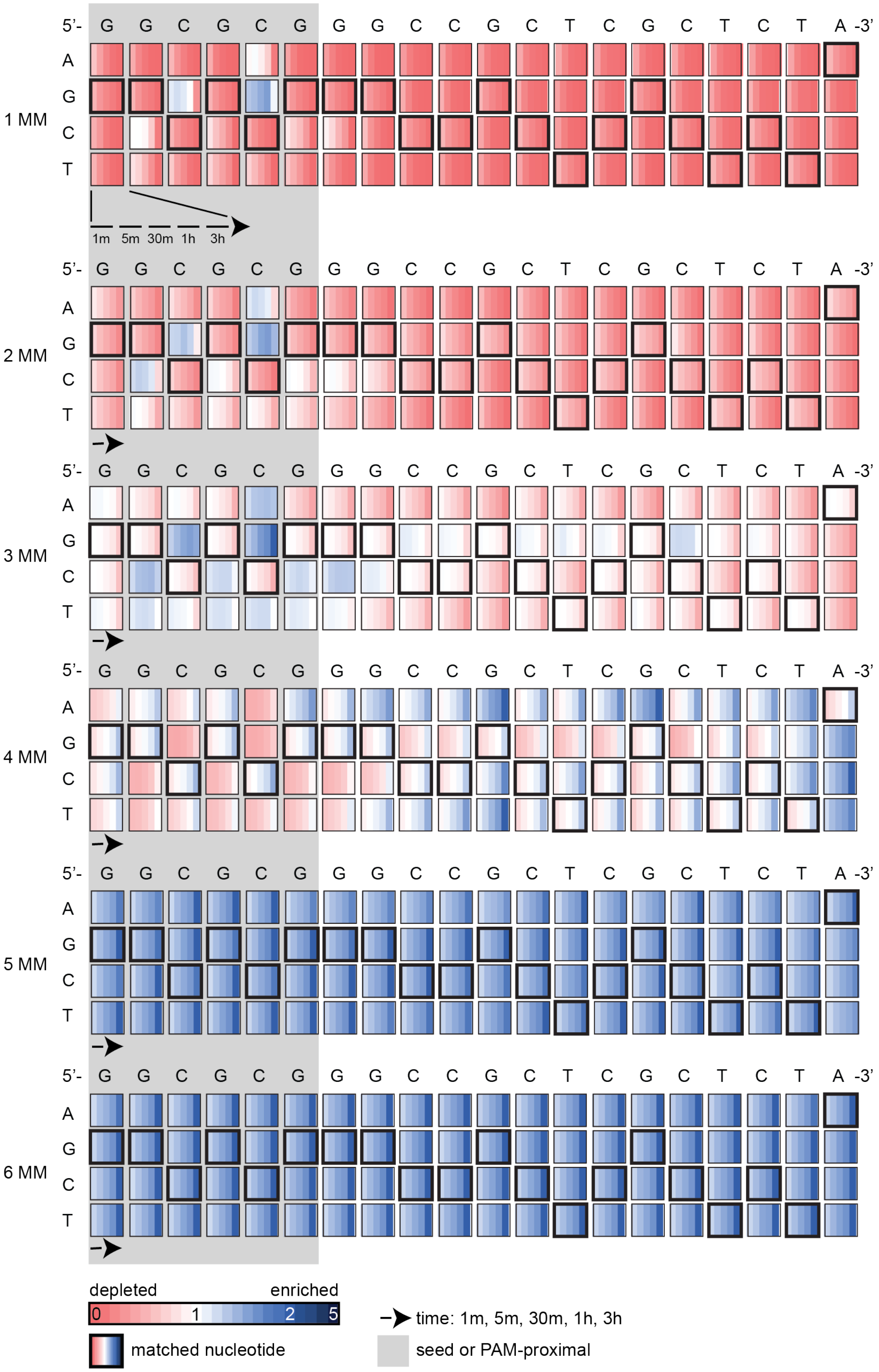
5'- G G C G C G G G C C G C T C G C T C T A -3'
A
G
C
T
1 MM
1m 5m 30m 1h 3h
2 MM
3 MM
4 MM
5 MM
6 MM
depleted
enriched
0
1
2
5
matched nucleotide
time: 1m, 5m, 30m, 1h, 3h
seed or PAM-proximal

**Supplementary Figure 8: Mismatched target sequences in the supercoiled fraction of pLibrary EMX1 upon cleavage by AsCas12a.**

Heatmaps showing the relative enrichment (blue) or depletion (red) of different mismatched sequences over time for the supercoiled fraction in pLibrary EMX1 upon cleavage by AsCas12a. The target sequence is indicated on the top and highlighted by bold black boxes in the heatmap. The PAM-proximal “seed” sequence is highlighted by the grey box. Each box represents the proportion of sequences containing each nucleotide at a given position across time. Values plotted represent average of three replicates. MM = mismatch

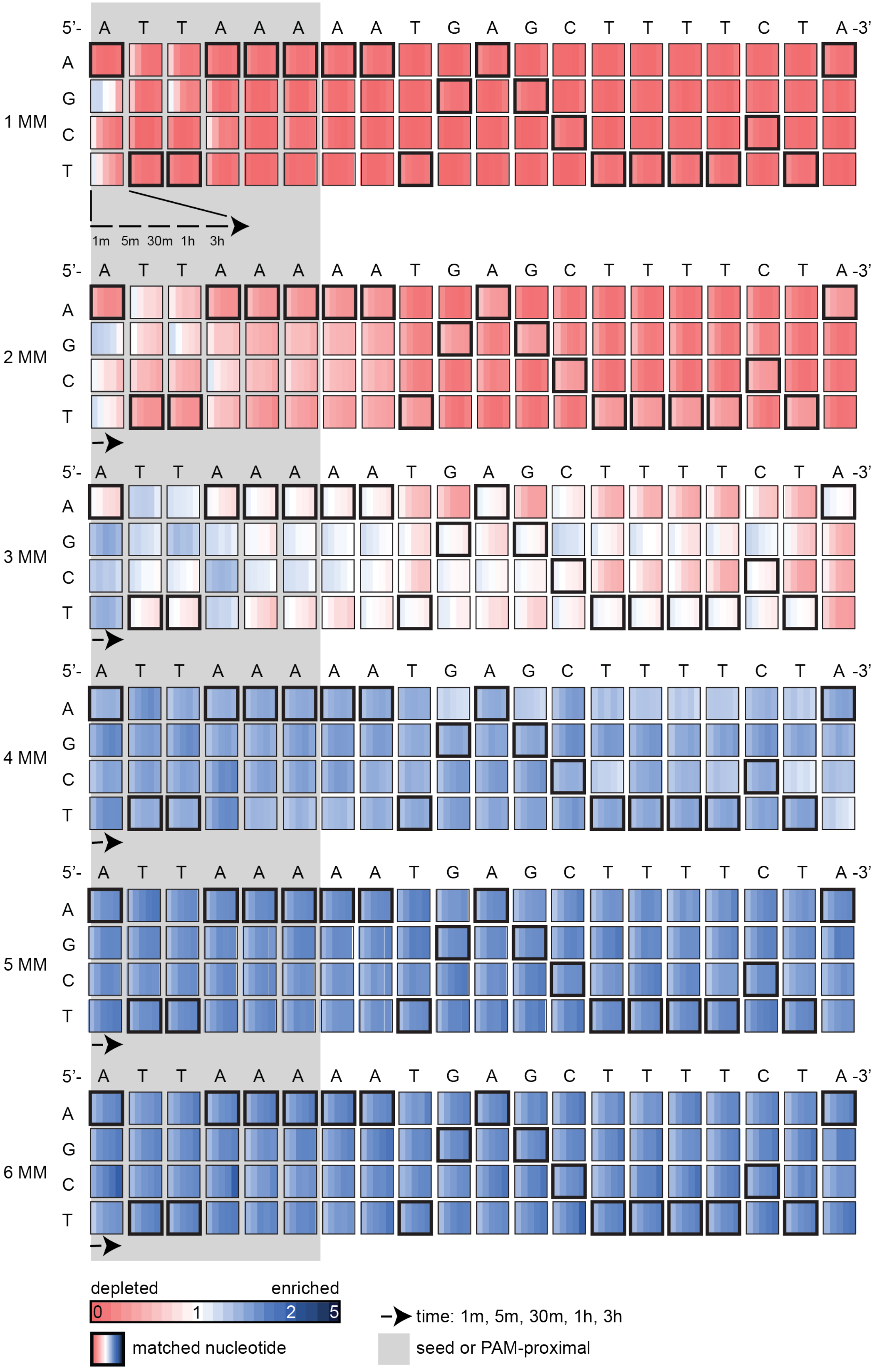

5'- A T T A A A A A T G A G C T T T T C T A -3'
A
G
C
T
1 MM
2 MM
3 MM
4 MM
5 MM
6 MM
1m 5m 30m 1h 3h
depleted
enriched
0
1
2
5
matched nucleotide
time: 1m, 5m, 30m, 1h, 3h
seed or PAM-proximal

**Supplementary Figure 9: Mismatched target sequences in the supercoiled fraction of pLibrary CCR5 upon cleavage by FnCas12a.**

Heatmaps showing the relative enrichment (blue) or depletion (red) of different mismatched sequences over time for the supercoiled fraction in pLibrary CCR5 upon cleavage by FnCas12a. The target sequence is indicated on the top and highlighted by bold black boxes in the heatmap. The PAM-proximal "seed" sequence is highlighted by the grey box. Each box represents the proportion of sequences containing each nucleotide at a given position across time. Values plotted represent average of three replicates. MM = mismatch

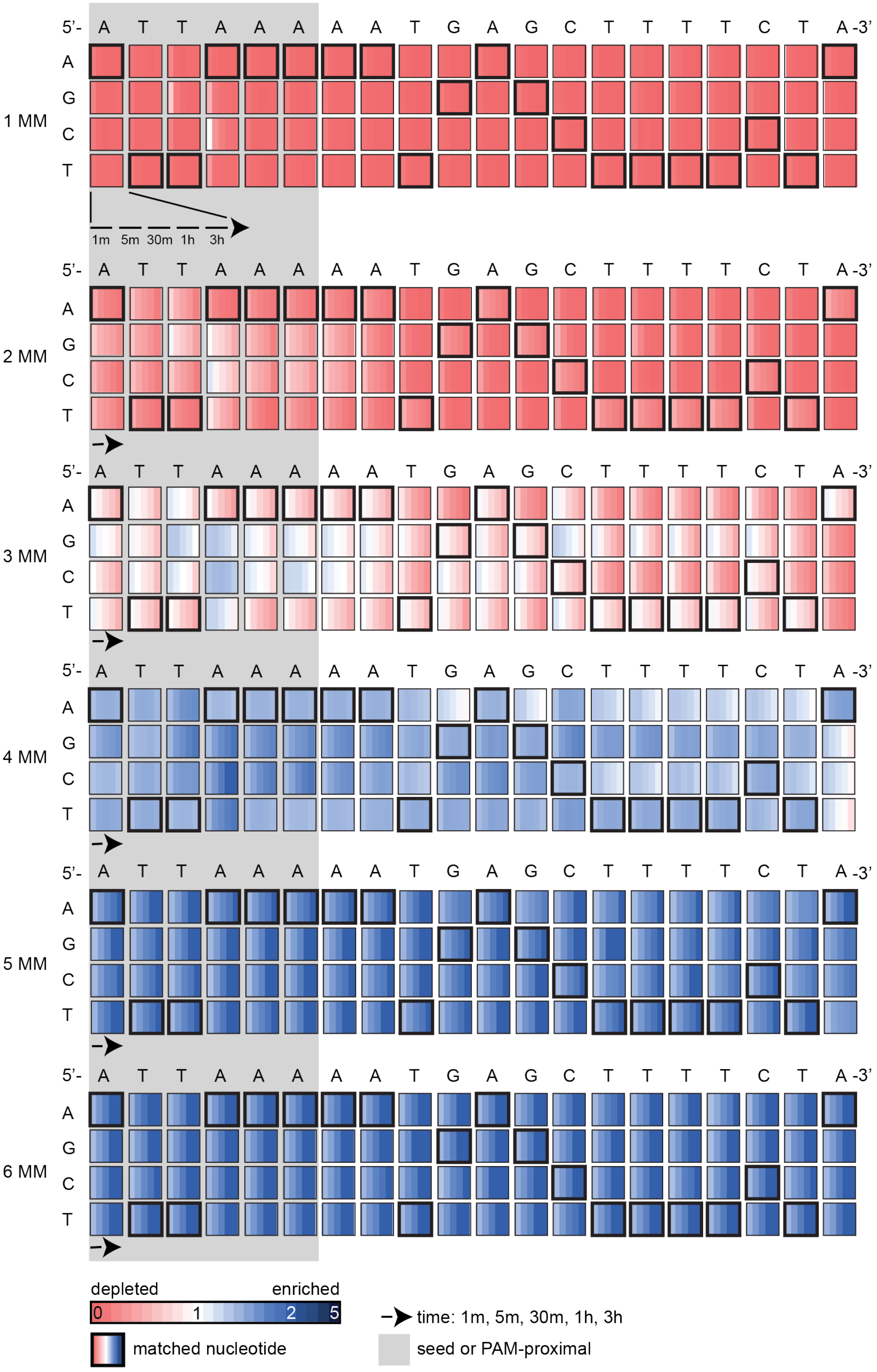
5'- A T T A A A A A T G A G C T T T T C T A -3'
A
G
C
T
1 MM
2 MM
3 MM
4 MM
5 MM
6 MM
1m 5m 30m 1h 3h
depleted
enriched
0
1
2
5
matched nucleotide
time: 1m, 5m, 30m, 1h, 3h
seed or PAM-proximal

**Supplementary Figure 10: Mismatched target sequences in the supercoiled fraction of pLibrary CCR5 upon cleavage by AsCas12a.**

Heatmaps showing the relative enrichment (blue) or depletion (red) of different mismatched sequences over time for the supercoiled fraction in pLibrary CCR5 upon cleavage by AsCas12a. The target sequence is indicated on the top and highlighted by bold black boxes in the heatmap. The PAM-proximal "seed" sequence is highlighted by the grey box. Each box represents the proportion of sequences containing each nucleotide at a given position across time. Values plotted represent average of three replicates. MM = mismatch

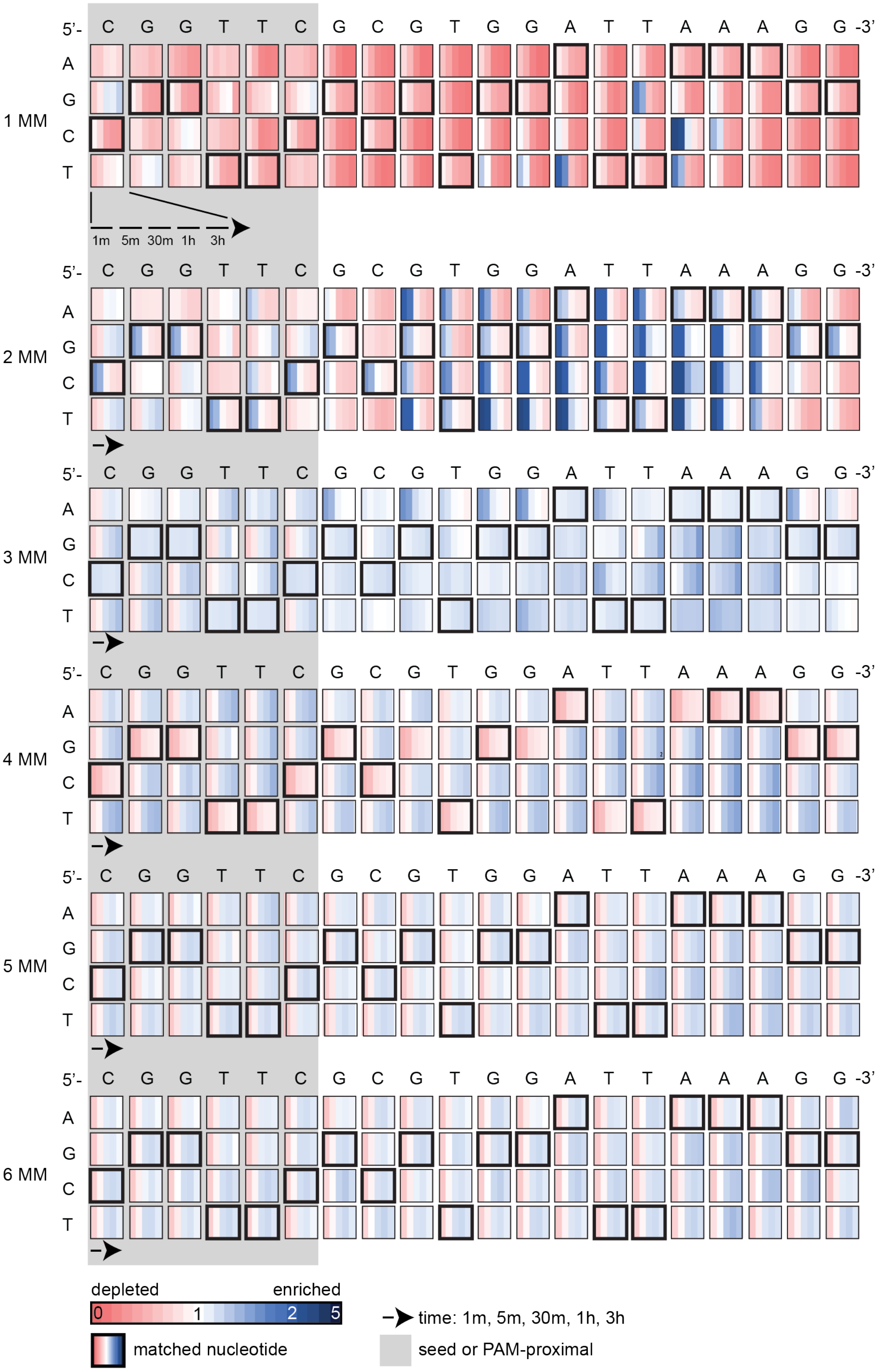

5'- C G G T T C G C G T G G A T T A A A G G -3'
A
G
C
T
1 MM
2 MM
3 MM
4 MM
5 MM
6 MM
1m 5m 30m 1h 3h
depleted
enriched
0
1
2
5
matched nucleotide
time: 1m, 5m, 30m, 1h, 3h
seed or PAM-proximal

**Supplementary Figure 11: Mismatched target sequences in the nicked fraction of pLibrary PS4 upon cleavage by FnCas12a.**

Heatmaps showing the relative enrichment (blue) or depletion (red) of different mismatched sequences over time for the nicked fraction in pLibrary PS4 upon cleavage by FnCas12a. The target sequence is indicated on the top and highlighted by bold black boxes in the heatmap. The PAM-proximal “seed” sequence is highlighted by the grey box. Each box represents the proportion of sequences containing each nucleotide at a given position across time. Values plotted represent average of three replicates. MM = mismatch

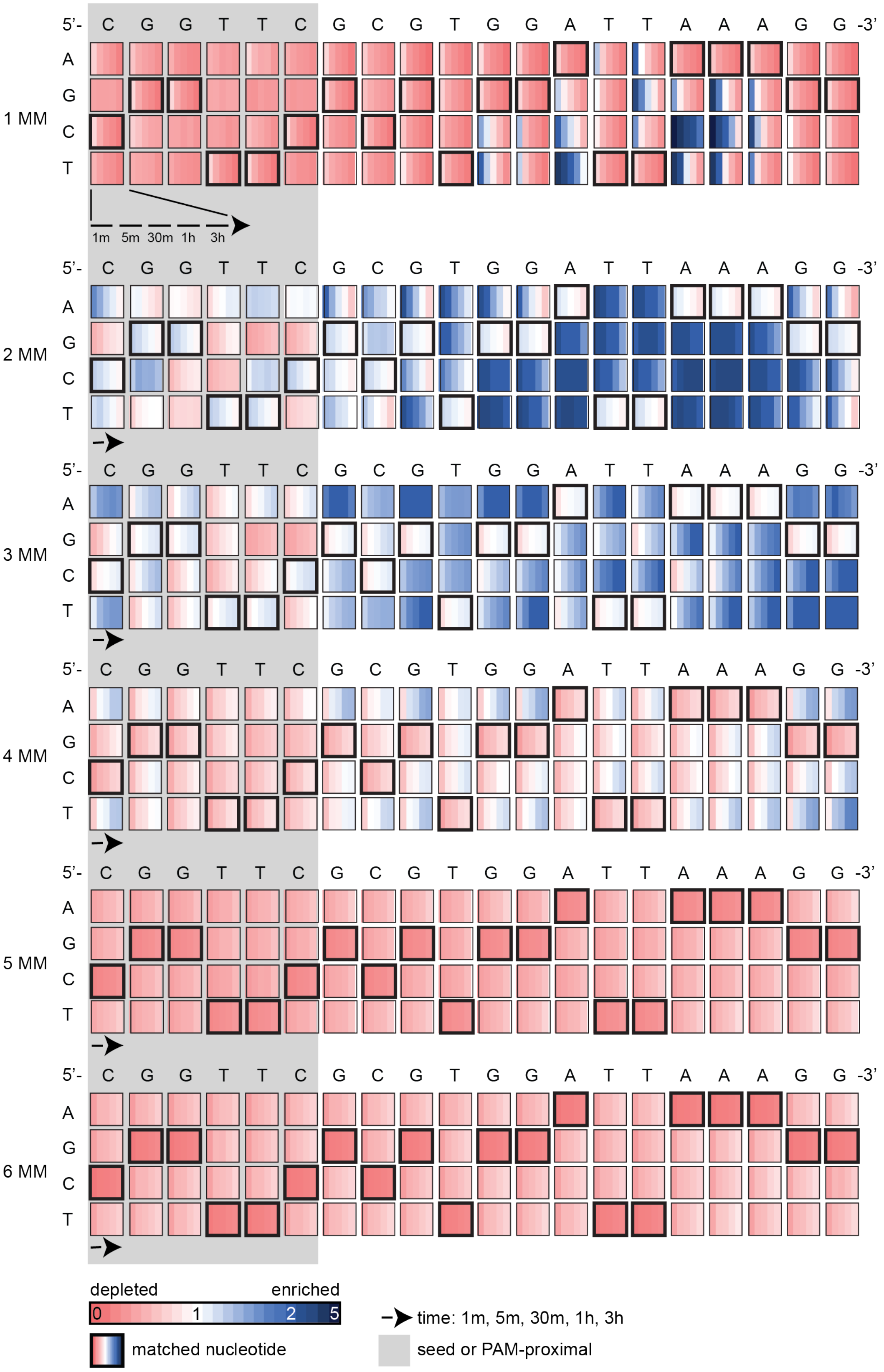

5'- C G G T T C G C G T G G A T T A A A G G -3'
A
G
C
T
1 MM
2 MM
3 MM
4 MM
5 MM
6 MM
1m 5m 30m 1h 3h
depleted
enriched
0
1
2
5
matched nucleotide
time: 1m, 5m, 30m, 1h, 3h
seed or PAM-proximal

**Supplementary Figure 12: Mismatched target sequences in the nicked fraction of pLibrary PS4 upon cleavage by AsCas12a.**

Heatmaps showing the relative enrichment (blue) or depletion (red) of different mismatched sequences over time for the nicked fraction in pLibrary PS4 upon cleavage by AsCas12a. The target sequence is indicated on the top and highlighted by bold black boxes in the heatmap. The PAM-proximal "seed" sequence is highlighted by the grey box. Each box represents the proportion of sequences containing each nucleotide at a given position across time. Values plotted represent average of three replicates. MM = mismatch

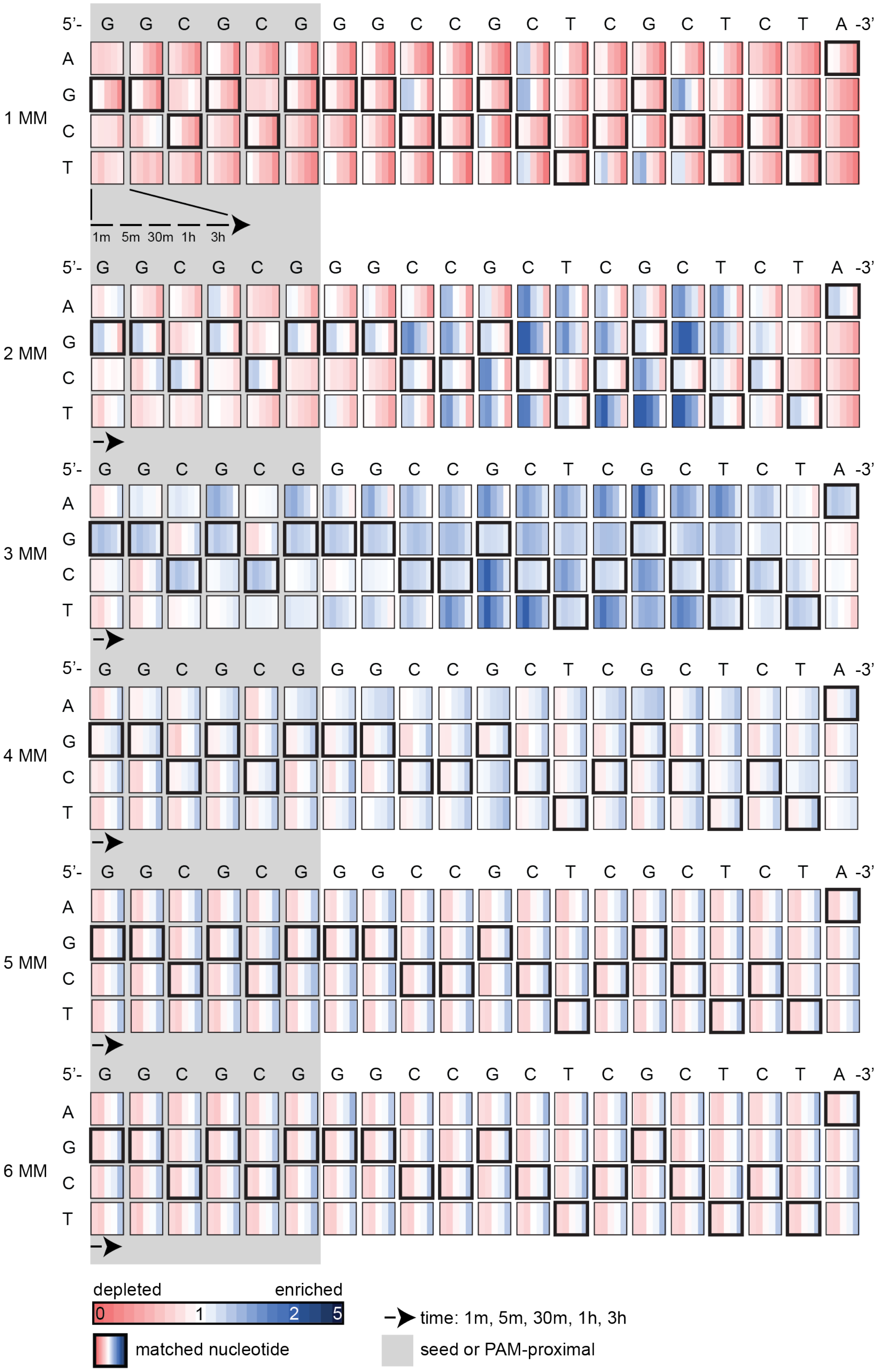

5'- G G C G C G G G C C G C T C G C T C T A -3'
A
G
C
T
1 MM
2 MM
3 MM
4 MM
5 MM
6 MM
1m 5m 30m 1h 3h
depleted
enriched
0
1
2
5
matched nucleotide
time: 1m, 5m, 30m, 1h, 3h
seed or PAM-proximal

**Supplementary Figure 13: Mismatched target sequences in the nicked fraction of pLibrary EMX1 upon cleavage by FnCas12a.**

Heatmaps showing the relative enrichment (blue) or depletion (red) of different mismatched sequences over time for the nicked fraction in pLibrary EMX1 upon cleavage by FnCas12a. The target sequence is indicated on the top and highlighted by bold black boxes in the heatmap. The PAM-proximal "seed" sequence is highlighted by the grey box. Each box represents the proportion of sequences containing each nucleotide at a given position across time. Values plotted represent average of three replicates. MM = mismatch

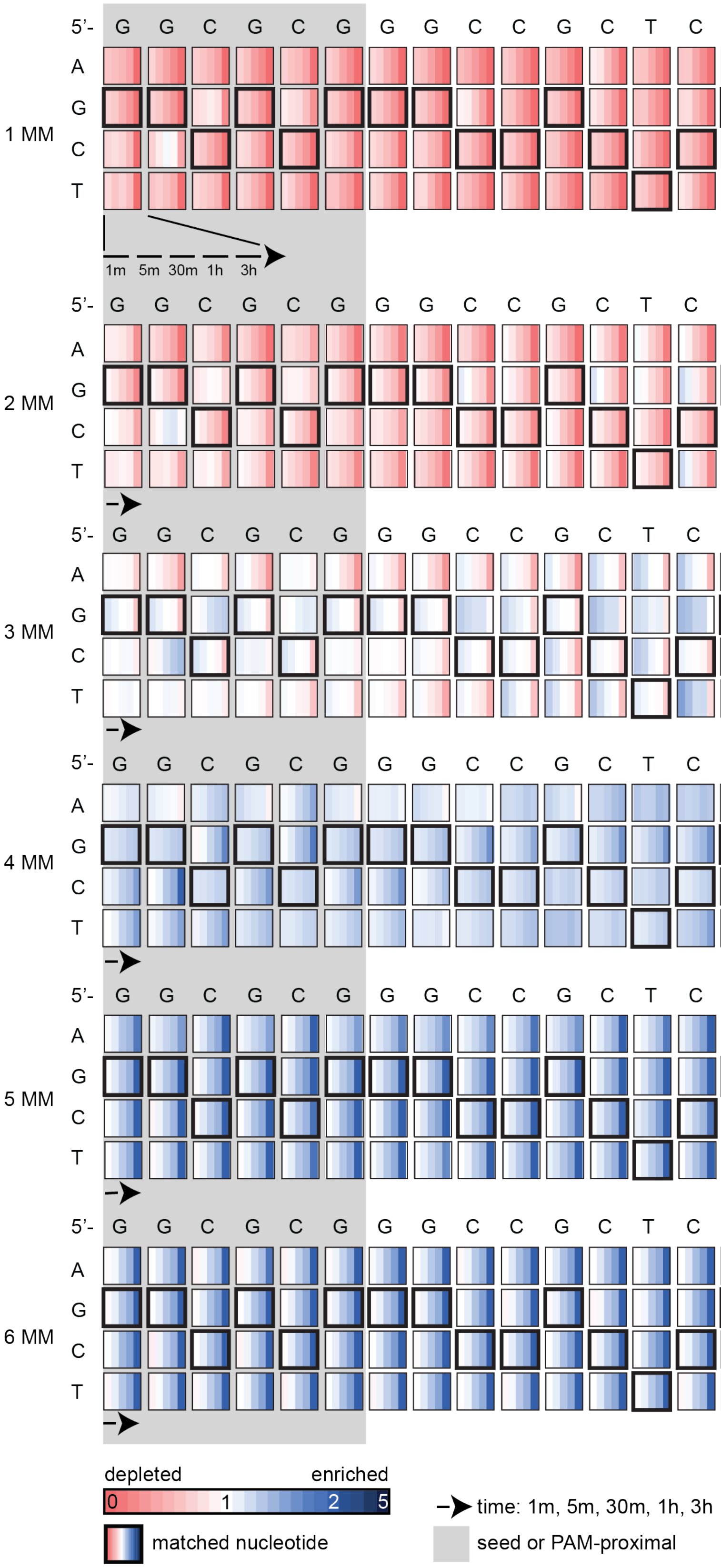

5'- G G C G C G G G C C G C T C G C T C T A -3'
A
G
C
T
1 MM
1m 5m 30m 1h 3h
2 MM
3 MM
4 MM
5 MM
6 MM
depleted
enriched
0
1
2
5
matched nucleotide
time: 1m, 5m, 30m, 1h, 3h
seed or PAM-proximal

**Supplementary Figure 14: Mismatched target sequences in the nicked fraction of pLibrary EMX1 upon cleavage by LbCas12a.**

Heatmaps showing the relative enrichment (blue) or depletion (red) of different mismatched sequences over time for the nicked fraction in pLibrary EMX1 upon cleavage by LbCas12a. The target sequence is indicated on the top and highlighted by bold black boxes in the heatmap. The PAM-proximal “seed” sequence is highlighted by the grey box. Each box represents the proportion of sequences containing each nucleotide at a given position across time. Values plotted represent average of three replicates. MM = mismatch

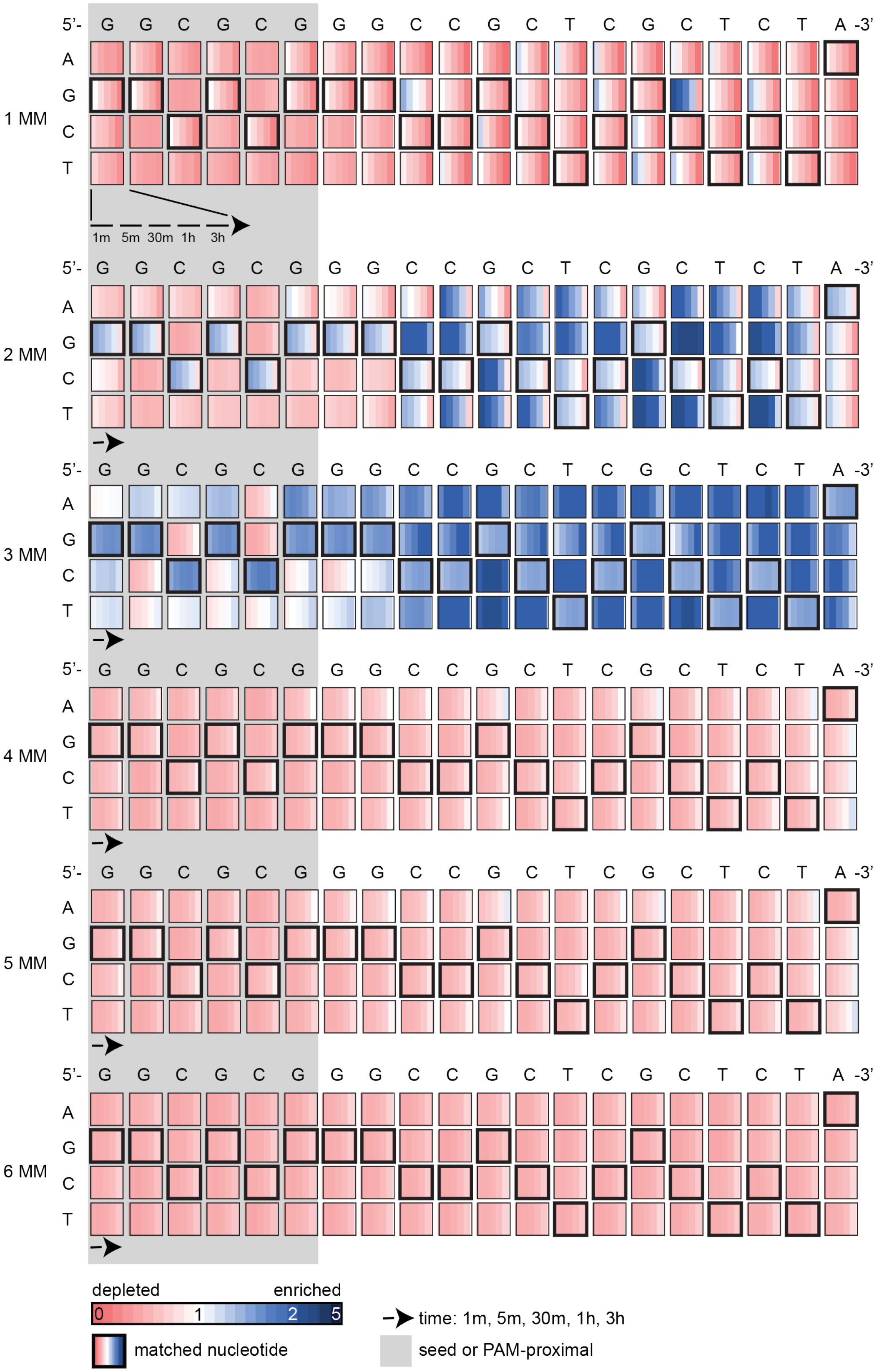
5'- G G C G C G G G C C G C T C G C T C T A -3'
A
G
C
T
1 MM
2 MM
3 MM
4 MM
5 MM
6 MM
1m 5m 30m 1h 3h
depleted
enriched
0
1
2
5
matched nucleotide
time: 1m, 5m, 30m, 1h, 3h
seed or PAM-proximal

**Supplementary Figure 15: Mismatched target sequences in the nicked fraction of pLibrary EMX1 upon cleavage by AsCas12a.**

Heatmaps showing the relative enrichment (blue) or depletion (red) of different mismatched sequences over time for the nicked fraction in pLibrary EMX1 upon cleavage by AsCas12a. The target sequence is indicated on the top and highlighted by bold black boxes in the heatmap. The PAM-proximal "seed" sequence is highlighted by the grey box. Each box represents the proportion of sequences containing each nucleotide at a given position across time. Values plotted represent average of three replicates. MM = mismatch

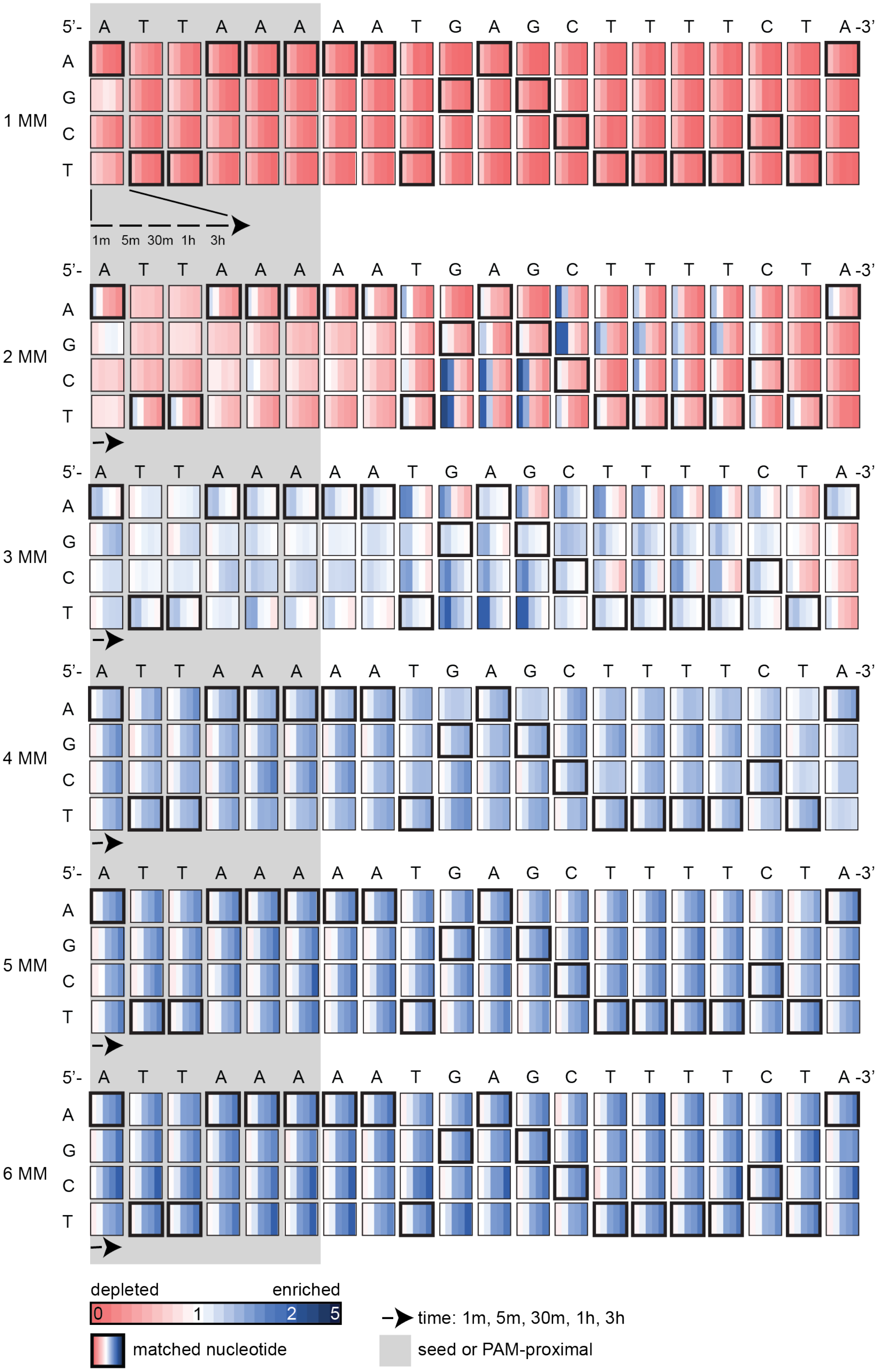

5'- A T T A A A A A T G A G C T T T T C T A -3'
A
G
C
T
1 MM
1m 5m 30m 1h 3h
2 MM
3 MM
4 MM
5 MM
6 MM
depleted
enriched
0
1
2
5
matched nucleotide
time: 1m, 5m, 30m, 1h, 3h
seed or PAM-proximal

**Supplementary Figure 16: Mismatched target sequences in the nicked fraction of pLibrary CCR5 upon cleavage by FnCas12a.**

Heatmaps showing the relative enrichment (blue) or depletion (red) of different mismatched sequences over time for the nicked fraction in pLibrary CCR5 upon cleavage by FnCas12a. The target sequence is indicated on the top and highlighted by bold black boxes in the heatmap. The PAM-proximal “seed” sequence is highlighted by the grey box. Each box represents the proportion of sequences containing each nucleotide at a given position across time. Values plotted represent average of three replicates. MM = mismatch

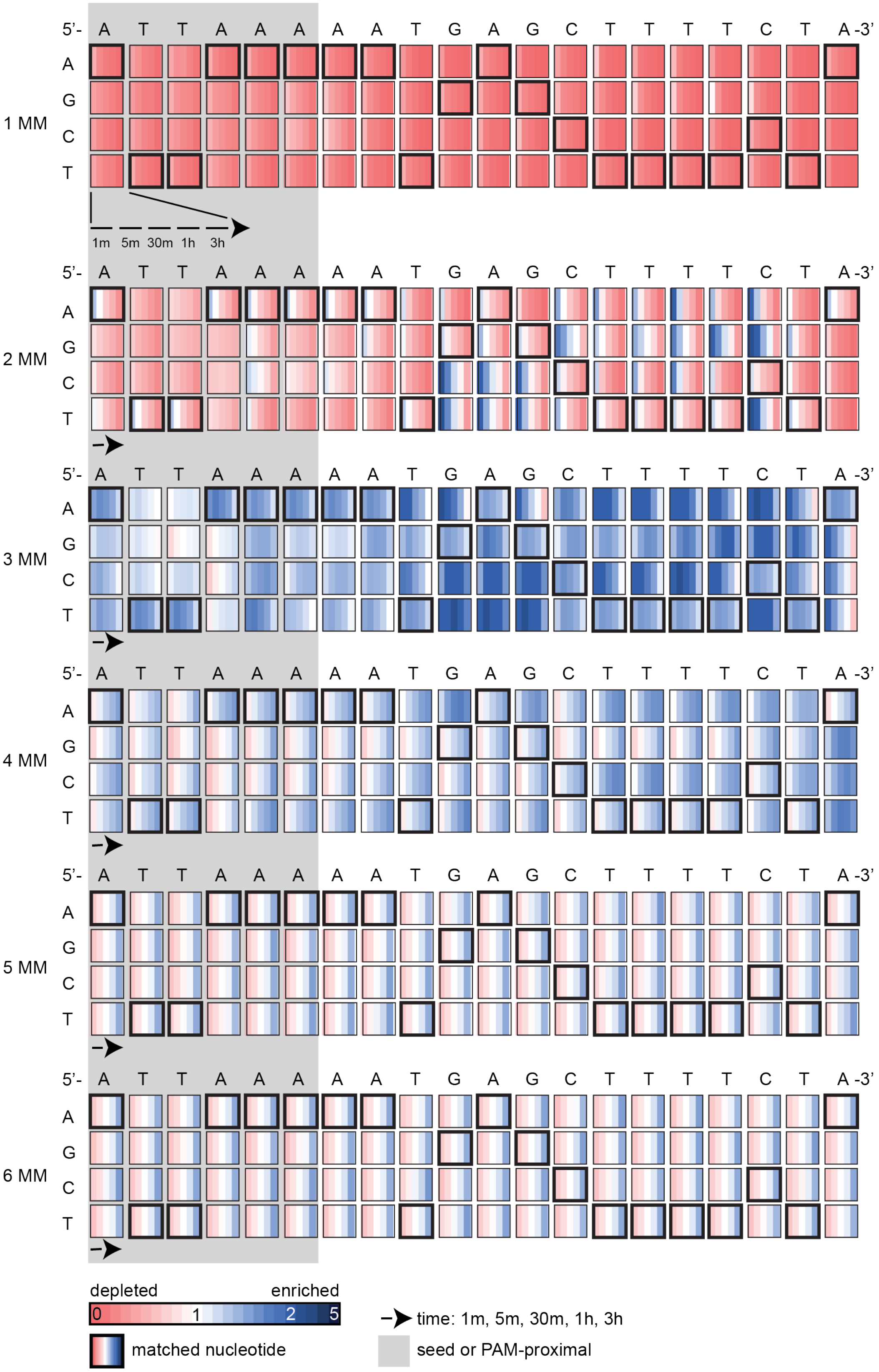

5'- A T T A A A A A T G A G C T T T T C T A -3'
A
G
C
T
1 MM
2 MM
3 MM
4 MM
5 MM
6 MM
1m 5m 30m 1h 3h
depleted
enriched
0 1 2 5
matched nucleotide
time: 1m, 5m, 30m, 1h, 3h
seed or PAM-proximal

**Supplementary Figure 17: Mismatched target sequences in the nicked fraction of pLibrary CCR5 upon cleavage by AsCas12a.**

Heatmaps showing the relative enrichment (blue) or depletion (red) of different mismatched sequences over time for the nicked fraction in pLibrary CCR5 upon cleavage by AsCas12a. The target sequence is indicated on the top and highlighted by bold black boxes in the heatmap. The PAM-proximal "seed" sequence is highlighted by the grey box. Each box represents the proportion of sequences containing each nucleotide at a given position across time. Values plotted represent average of three replicates. MM = mismatch

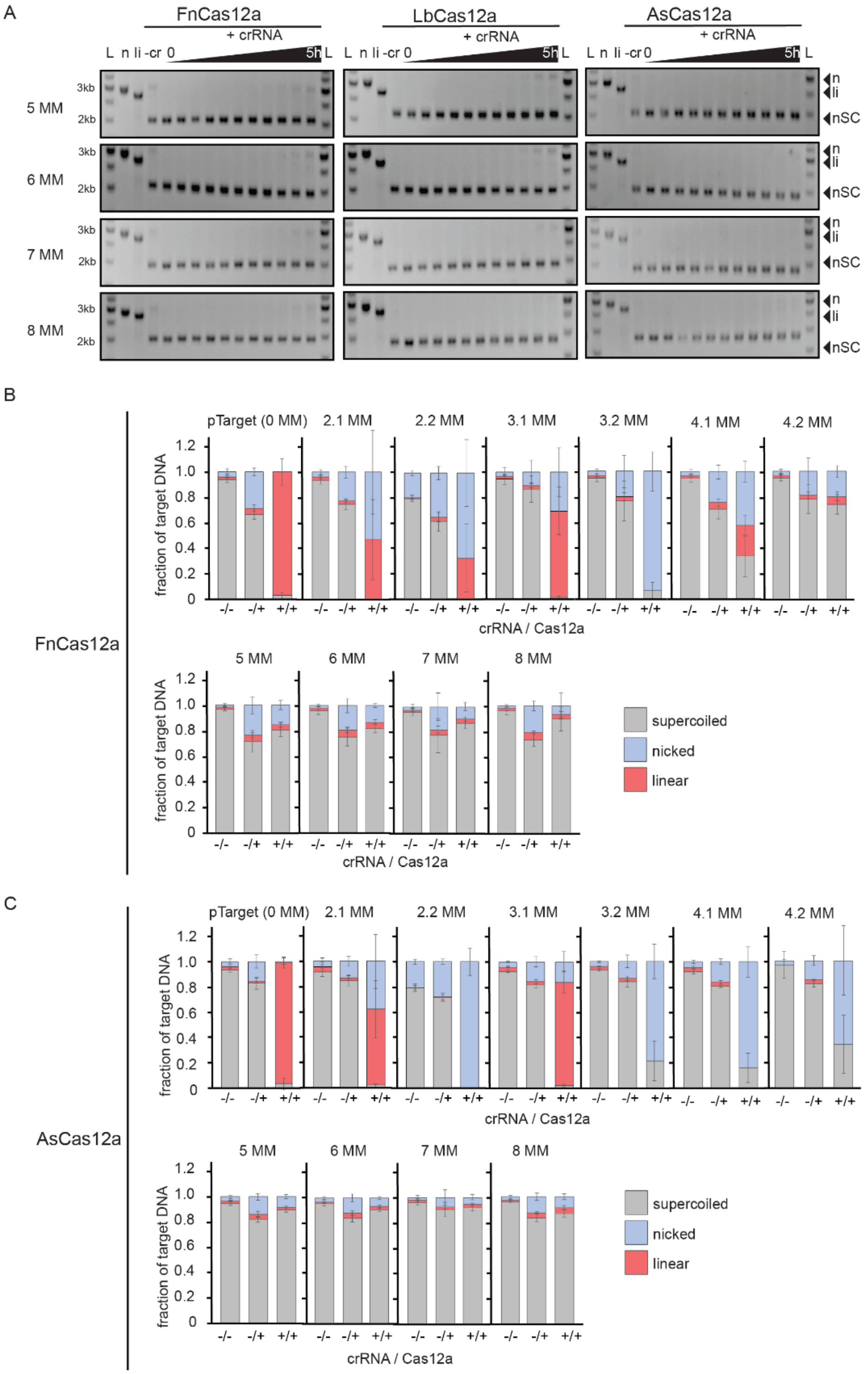


**Supplementary Figure 18: Cas12a orthologs cannot cleave target sequences with five or more mismatches.**

(A) Representative agarose gels showing no significant cleavage of a negatively supercoiled (nSC) plasmid containing the mismatched (MM) target over a time course by Cas12a orthologs,

FnCas12a (left), LbCas12a (center) and AsCas12a (right). Time points at which the samples were collected are 15 sec, 30 sec, 1 min, 2 min, 5 min, 15 min, 30 min, 1 hour, 3 hours, and 5 hours.

Controls: -cr = reaction without cognate crRNA, n = Nt.BspQI nicked pUC19, li = BsaI-HF linearized pUC19.

(B – C) Quantification of supercoiled, linear and nicked fractions from cleavage of perfect and mismatched (MM) target plasmid by (B) FnCas12a and (C) AsCas12a after 3 hours. -/- indicates a cleavage reaction with the target plasmid without Cas12a and crRNA. -/+ indicates a cleavage reaction with the target plasmid and Cas12a only, and +/+ indicates a cleavage reaction with the target plasmid, Cas12a and cognate crRNA. Average of the intensity fraction values are plotted with SD as error bars, n = 3 replicates.

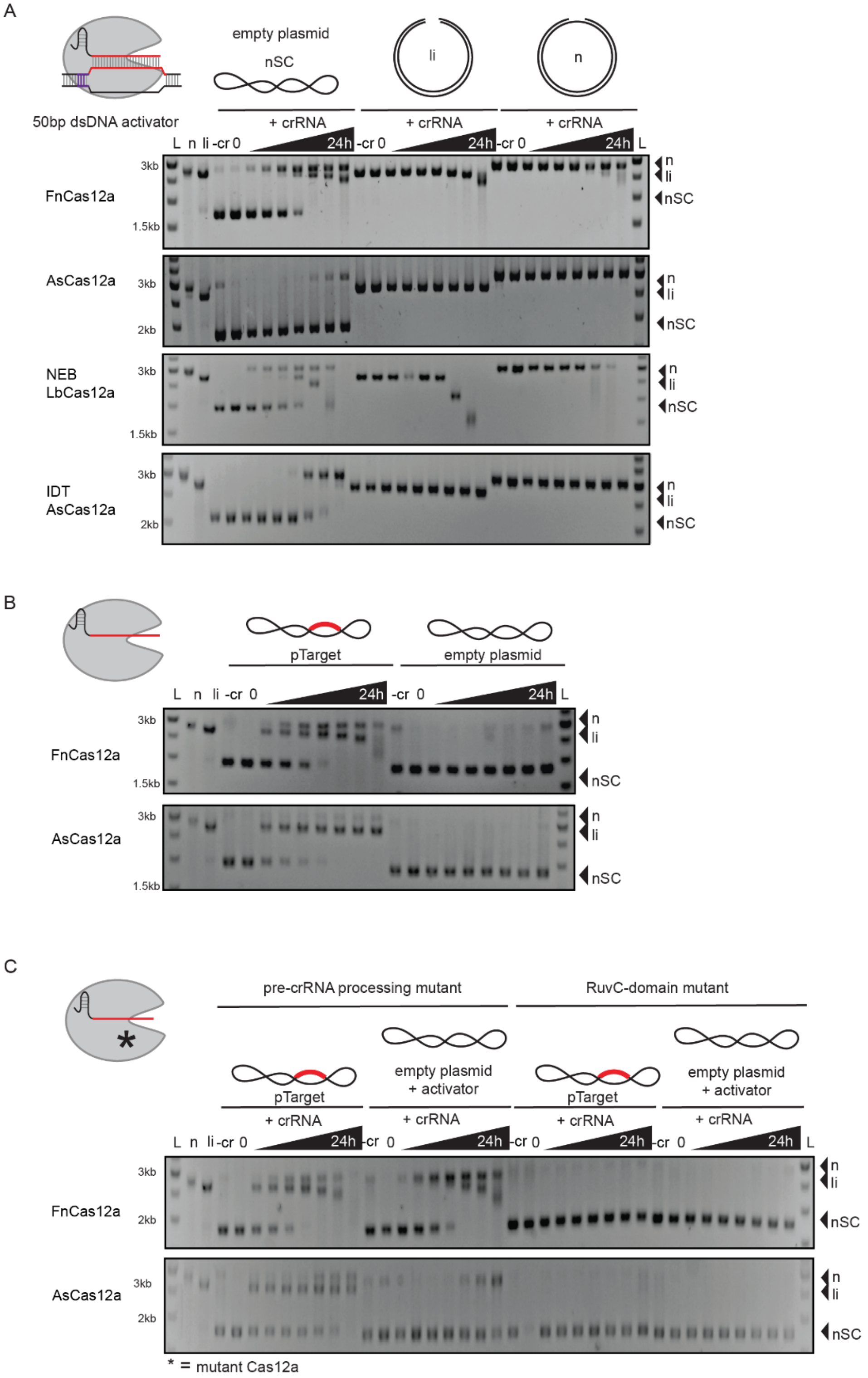


**Supplementary Figure 19: FnCas12a and LbCas12a have strong activated non-specific, trans dsDNA nicking and degradation activity.**

(A) Representative agarose gel showing non-specific nicking and linearization of negatively supercoiled (nSC) dsDNA plasmid, and degradation of linearized and nicked dsDNA plasmid over time by Cas12a orthologs (lab-made and commercially available). Cas12a (20 nM) and crRNA (30 nM) were complexed with a crRNA-complementary 50 bp dsDNA activator (30 nM) (perfect target from pLibrary PS4) and incubated with non-specific dsDNA in different forms – negatively supercoiled (nSC), linear (li) and nicked (n).
(B) Representative agarose gel showing target-dependent cleavage and degradation of dsDNA by FnCas12a and LbCas12a.
(C) Representative agarose gel showing RuvC-domain dependent pTarget linearization and activated non-specific nicking and degradation of dsDNA plasmid. Pre-crRNA processing and RuvC-domain active site mutants of Cas12a (20 nM) and crRNA (30 nM) were complexed with a crRNA-complementary 50bp dsDNA activator (30 nM) (perfect target from pLibrary PS4) and incubated with non-specific dsDNA in different forms – negatively supercoiled (nSC), linear (li) and nicked (n).
Time points at which the samples were collected are 5 min, 15 min, 30 min, 1 hour, 4 hours, 8 hours, and 24 hours. Controls: -cr = reaction without cognate crRNA, n = Nt.BspQI nicked pUC19, li = BsaI-HF linearized pUC19.

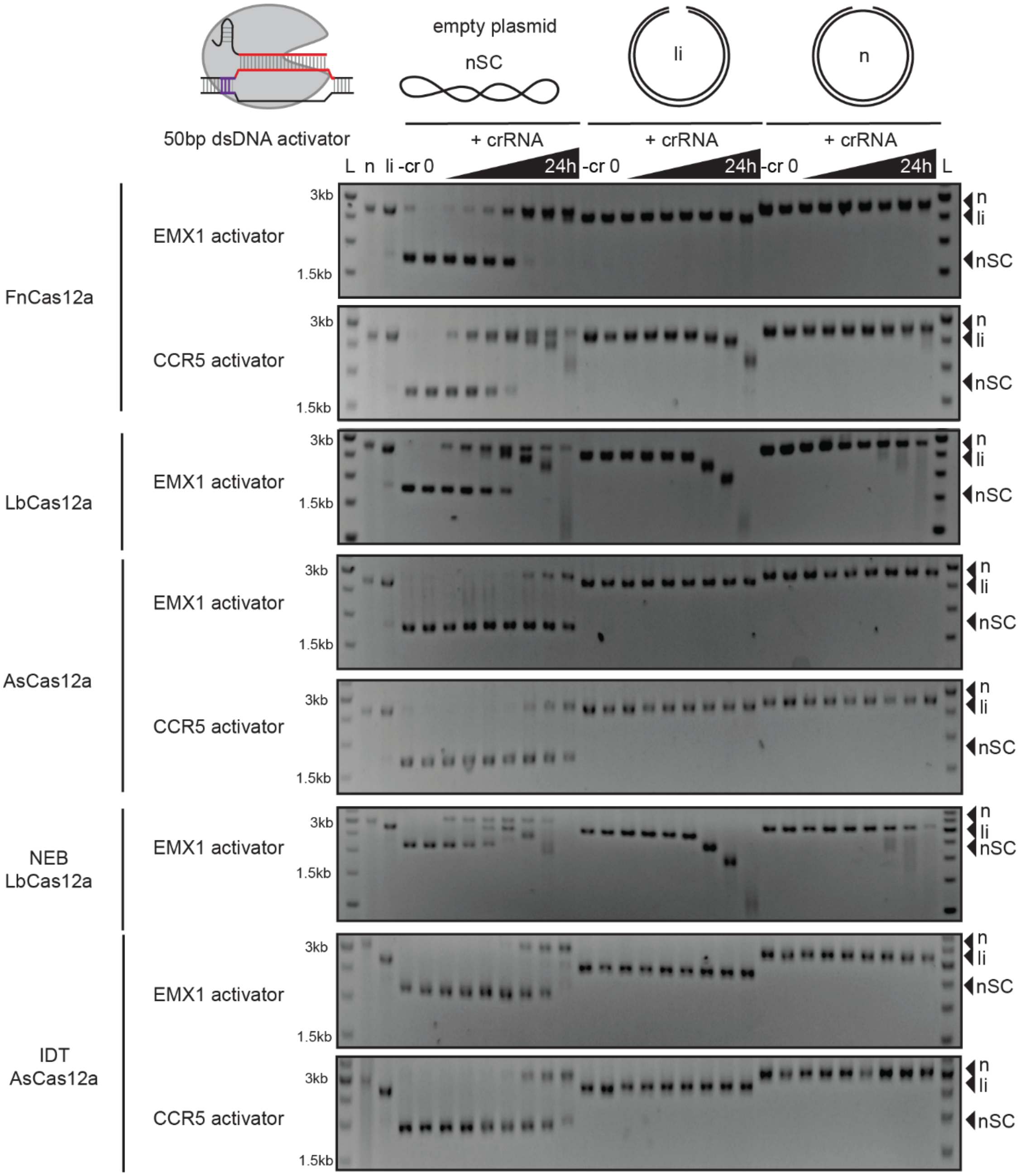


**Supplementary Figure 20: Cas12a can be activated for non-specific, trans dsDNA nicking and degradation activity by different crRNA-activator pairs.**

Representative agarose gels showing non-specific nicking and linearization of negatively supercoiled (nSC) dsDNA plasmid, and degradation of linearized and nicked dsDNA plasmid by different Cas12a orthologs (lab-made and commercially available). Cas12a (20 nM) and crRNA (30 nM) were complexed with a crRNA-complementary 50bp dsDNA activator (30 nM) (as indicated in the figure) and incubated with non-specific dsDNA in different forms – negatively supercoiled (nSC), linear (li) and nicked (n). Different crRNA-activator pairs were tested against non-specific dsDNA. Time points at which the samples were collected are 5 min, 15 min, 30 min, 1 hour, 4 hours, 8 hours, and 24 hours.

Controls: -cr = reaction without cognate crRNA, n = Nt.BspQI nicked pUC19, li = BsaI-HF linearized pUC19.

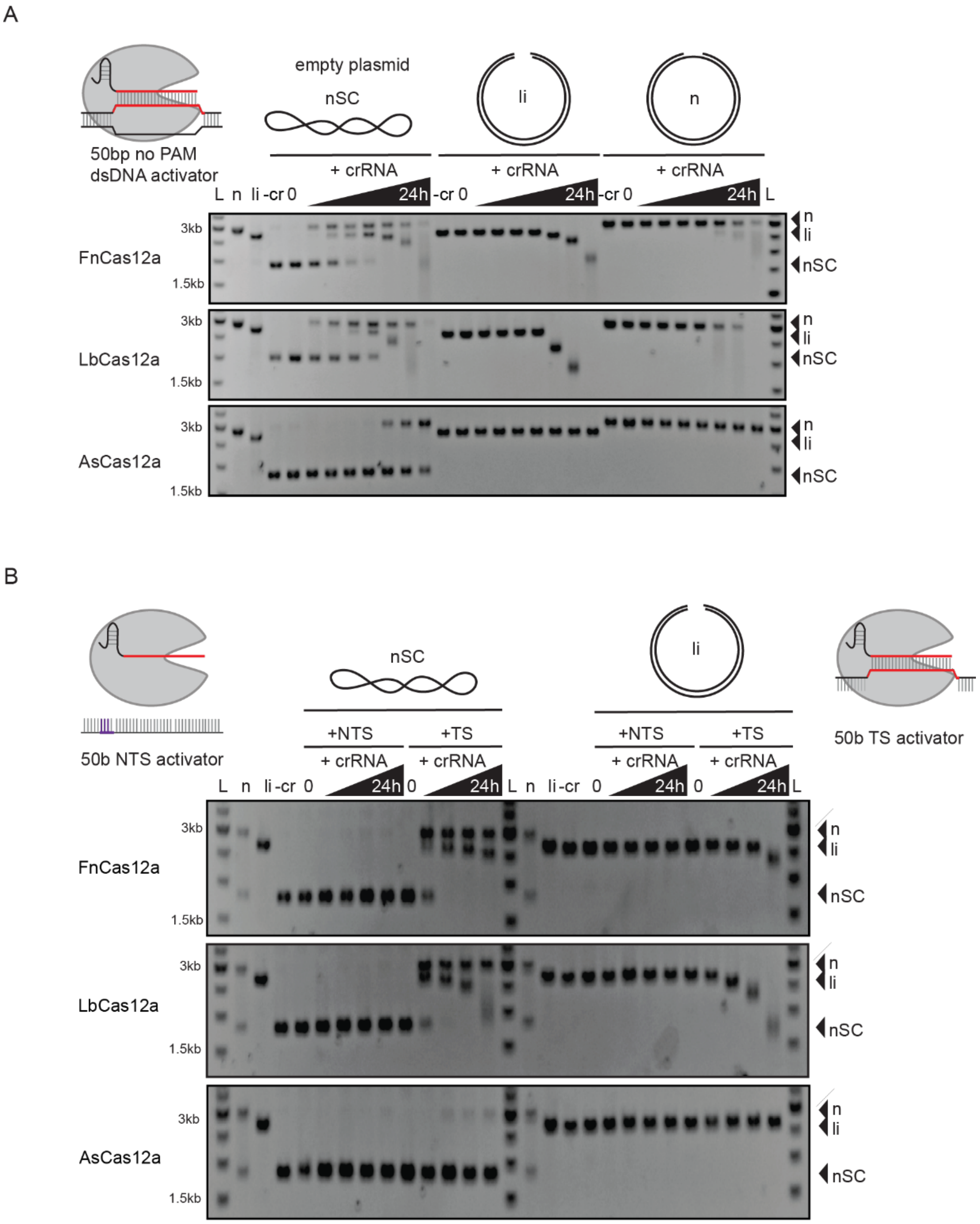


**Supplementary Figure 21: Requirements for activation of Cas12a for trans dsDNA nicking and degradation activities.**

(A) Representative agarose gels showing non-specific nicking and linearization of negatively supercoiled (nSC) dsDNA plasmid, and degradation of linearized dsDNA plasmid by Cas12a orthologs upon activation in the absence of PAM. Cas12a (20 nM) and crRNA (30 nM) were complexed with a 50bp no PAM dsDNA activator (30 nM) and incubated with non-specific dsDNA in different forms – negatively supercoiled (nSC), linear (li) and nicked (n). Time points at which the samples were collected are 5 min, 15 min, 30 min, 1 hour, 4 hours, 8 hours, and 24 hours.

(B) Representative agarose gels showing non-specific nicking and linearization of nSC dsDNA plasmid, and degradation of linearized dsDNA plasmid by Cas12a orthologs upon activation by only target-strand (TS) binding. Cas12a (20 nM) and crRNA (30 nM) were complexed with a 50bp ssDNA target strand (TS) or non-target strand (NTS) activator (30 nM) and incubated with non-specific negatively supercoiled (nSC) and linear (li) dsDNA. Time points at which the samples were collected are 1 hour, 4 hours, 8 hours, and 24 hours.

Controls: -cr = reaction without cognate crRNA, n = Nt.BspQI nicked pUC19, li = BsaI-HF linearized pUC19.

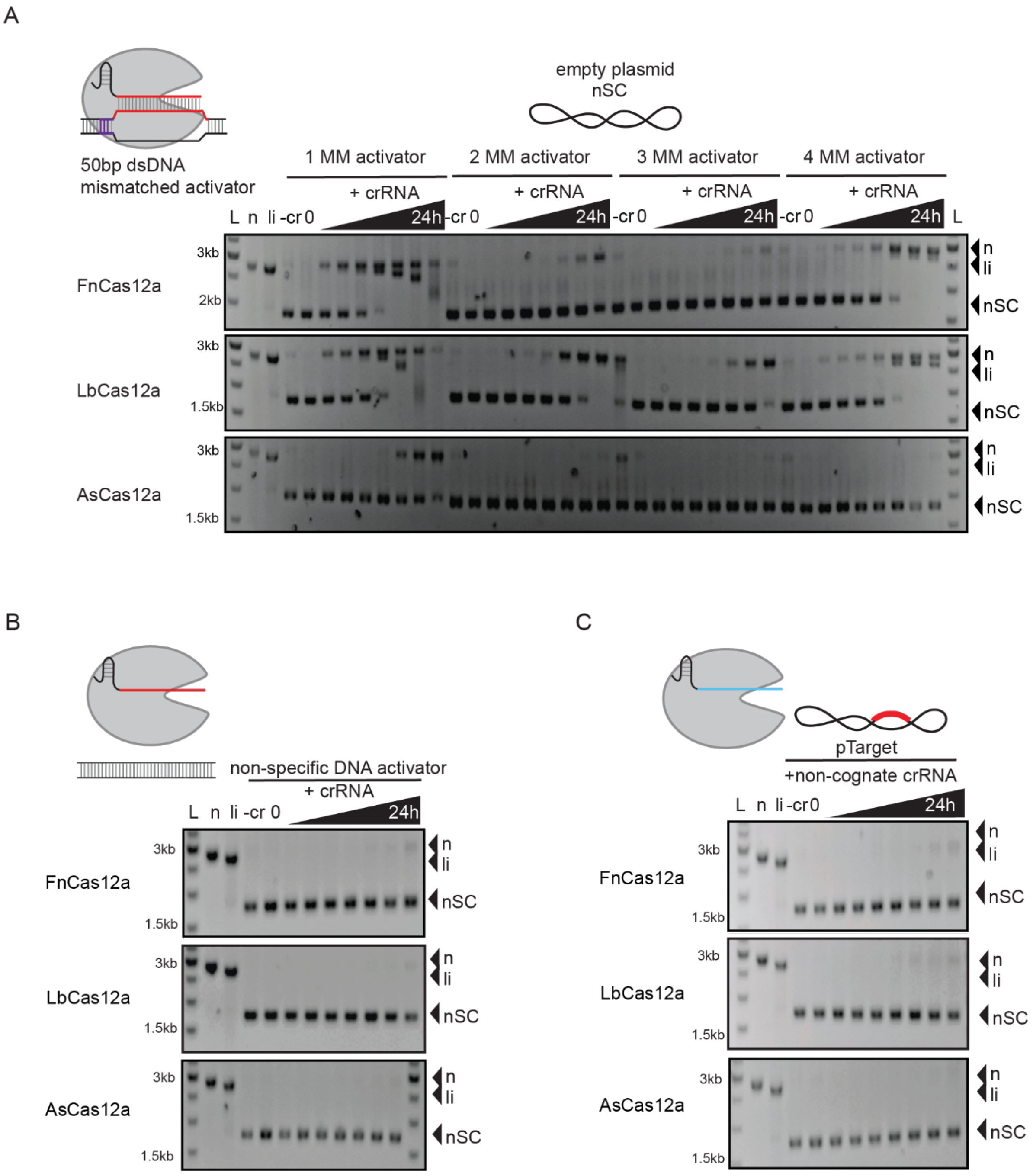


**Supplementary Figure 22: Cas12a can be activated for non-specific, trans dsDNA nicking and degradation activity by activators with up to four mismatches.**

(A) Representative agarose gels showing non-specific nicking and linearization of negatively supercoiled (nSC) dsDNA plasmid, and degradation of linearized dsDNA plasmid by Cas12a orthologs upon activation by target sequences with mismatches. Cas12a (10 nM) and crRNA (30 nM) were complexed with a 50bp dsDNA activator (30 nM) containing 1 to 4 mismatches (MM)

and incubated with non-specific negatively supercoiled (nSC) dsDNA.
(B – C) Representative agarose gels showing no cleavage of plasmid by Cas12a in the absence of (B) a specific activator or (C) cognate-crRNA. Cas12a (20 nM) and crRNA (30 nM) were complexed with a 50bp non-specific dsDNA activator (30 nM) and incubated with non-specific negatively supercoiled (nSC) dsDNA (left). Cas12a and a non-cognate crRNA were complexed and incubated with target containing dsDNA plasmid (right).
Time points at which the samples were collected are 5 min, 15 min, 30 min, 1 hour, 4 hours, 8 hours, and 24 hours. Controls: -cr = reaction without cognate crRNA, n = Nt.BspQI nicked pUC19, li = BsaI-HF linearized pUC19.

**Supplementary Table 1: List of Oligonucleotides**

Key:

**Bold** = spacer or target sequence

underlined = PAM

lowercase = mismatch (MM)

RC = reverse complement

| Sequence (5' to 3') | Notes |
|---|---|
| **RNA** | |
| AAUUUCUACUGUUGUAGAU**CGGUUCGCGUGGAUUAAAGG** | FnCas12a crRNA for modified protospacer 4 from Sp CRISPR |
| AAUUUCUACUAAGUGUAGAU**CGGUUCGCGUGGAUUAAAGG** | LbCas12a crRNA for modified protospacer 4 from Sp CRISPR |
| UAAUUUCUACUCUUGUAGAU**CGGUUCGCGUGGAUUAAAGG** | AsCas12a crRNA for modified protospacer 4 from Sp CRISPR |
| AAUUUCUACUGUUGUAGAU**GGCGCGGGCCGCUCGCUCUA** | FnCas12a - crRNA for EMX1 gene target |
| AAUUUCUACUAAGUGUAGAU**GGCGCGGGCCGCUCGCUCUA** | LbCas12a - crRNA for EMX1 gene target |
| UAAUUUCUACUCUUGUAGAU**GGCGCGGGCCGCUCGCUCUA** | AsCas12a - crRNA for EMX1 gene target |
| AAUUUCUACUGUUGUAGAU**AUUAAAAAUGAGCUUUUCUA** | FnCas12a crRNA for CCR5 gene target |
| AAUUUCUACUAAGUGUAGAU**AUUAAAAAUGAGCUUUUCUA** | LnCas12a crRNA for CCR5 gene target |
| UAAUUUCUACUCUUGUAGAU**AUUAAAAAUGAGCUUUUCUA** | AsCas12a crRNA for CCR5 gene target |
| **Target DNA oligonucleotides** | |
| GCATTGCTGTACGAATCGTACAGGGTGCTTCAGGATGTTTT**CGGTTCGCGTGGATTAAAGG**TGCGTCAAGCTCGGACATCGTGATTGATAATGCGATGC | Cas12a modified ps4 - DT1 - 99b target - ssoligo used for Gibson |

| | |
|---|---|
| | assembly with pUC19 – KMlib001 – pLibrary PS4 |
| GCATTGCTGTACGAATCGTACAGGGTGCTTCAGGTTTA**GGCGCGGGCCGCTCGCTCTA**GGGGGTGTCAAGCTCGGACATCGTGATTGATAATGCGATGC | EMX1 gene target - high GC % - 99b target - ssoligo used for Gibson assembly with pUC19 - KMlib003 – pLibrary EMX1 |
| GCATTGCTGTACGAATCGTACAGGGTGCTTCAGGTTTA**ATTAAAAATGAGCTTTTCTA**GGGGGTGTCAAGCTCGGACATCGTGATTGATAATGCGATGC | CCR5 gene target - low GC % - 99b target - ssoligo used for Gibson assembly with pUC19 - KMlib004 – pLibrary CCR5 |
| GCATTGCTGTACGAATCGTACAGGGTGCTTCAGGATGTTTA**CGGTTCGCtTGGATTtAAGG**TGCGTCAAGCTCGGACATCGTGATTGATAATGCGATGC | Cas12a mismatched target ssoligo - TTTA PAM - off target for mod protospacer 4, PLibrary PS4 - 2 mismatches 1 (2.1 MM) |
| GCATTGCTGTACGAATCGTACAGGGTGCTTCAGGATGTTTA**CGGTTCGCGTGGtTTAtAGG**TGCGTCAAGCTCGGACATCGTGATTGATAATGCGATGC | Cas12a mismatched target ssoligo - TTTA PAM - off target for mod protospacer 4, PLibrary PS4 - 2 mismatches 1 (2.2 MM) |
| GCATTGCTGTACGAATCGTACAGGGTGCTTCAGGATGTTTA**CGGTTCaCaTGGATTAgAGG**TGCGTCAAGCTCGGACATCGTGATTGATAATGCGATGC | Cas12a mismatched target ssoligo - TTTA PAM - off target for mod protospacer 4, PLibrary PS4 - 3 mismatches 1 (3.1 MM) |
| GCATTGCTGTACGAATCGTACAGGGTGCTTCAGGATGTTTA**CGGTcCGCGTtGATTAgAGG**TGCGTCAAGCTCGGACATCGTGATTGATAATGCGATGC | Cas12a mismatched target ssoligo - TTTA PAM - off target for mod protospacer 4, PLibrary PS4 – 3 mismatches 2 (3.2 MM) |
| GCATTGCTGTACGAATCGTACAGGGTGCTTCAGGATGTTTA**CacTTCGCGTGGATTtAAaG**TGCGTCAAGCTCGGACATCGTGATTGATAATGCGATGC | Cas12a mismatched target ssoligo - TTTA PAM - off target for mod protospacer 4, PLibrary PS4 – 4 mismatches 1 (4.1 MM) |

| | |
|---|---|
| GCATTGCTGTACGAATCGTACAGGGTGCTTCAGGATGTTTA**gGGTTCGtcTGGATTAgAGG**TGCGTCAAGCTCGGACATCGTGATTGATAATGCGATGC | Cas12a mismatched target ssoligo - TTTA PAM - off target for mod protospacer 4, PLibrary PS4 – 4 mismatches 2 (4.2 MM) |
| GCATTGCTGTACGAATCGTACAGGGTGCTTCAGGATGTTTA**aGGTaCGCGTGGgaTAtAGG**TGCGTCAAGCTCGGACATCGTGATTGATAATGCGATGC | Cas12a mismatched target ssoligo - TTTA PAM - off target for mod protospacer 4, PLibrary PS4 - 5 mismatches (5 MM) |
| GCATTGCTGTACGAATCGTACAGGGTGCTTCAGGATGTTTA**CtGTTCGaGTaGATTtAtGc**TGCGTCAAGCTCGGACATCGTGATTGATAATGCGATGC | Cas12a mismatched target ssoligo - TTTA PAM - off target for mod protospacer 4, PLibrary PS4 - 6 mismatches (6 MM) |
| GCATTGCTGTACGAATCGTACAGGGTGCTTCAGGATGTTTA**CGGtTtgCGTGGATTctttG**TGCGTCAAGCTCGGACATCGTGATTGATAATGCGATGC | Cas12a mismatched target ssoligo - TTTA PAM - off target for mod protospacer 4, PLibrary PS4 - 7 mismatches (7 MM) |
| GCATTGCTGTACGAATCGTACAGGGTGCTTCAGGATGTTTA**CGcggCtCGTtGtTTAcAtG**TGCGTCAAGCTCGGACATCGTGATTGATAATGCGATGC | Cas12a mismatched target ssoligo - TTTA PAM - off target for mod protospacer 4, PLibrary PS4 - 8 mismatches (8 MM) |
| **DNA oligonucleotide activators** | |
| GTGCTTCAGGATGTTTA**CGGTTCGCGTGGATTAAAGG**TGCGTCAAGCTCG | Cas12a modified ps4 - DT1 - 50b target |
| CGAGCTTGACGCACCTTTAATCCACGCGAACCGTAAACATCCTGAAGCAC | Cas12a modified ps4 - DT1 - 50b target - RC |
| GTGCTTCAGGATGTTTA**GGCGCGGGCCGCTCGCTCTA**TGCGTCAAGCTCG | Cas12a EMX1 - DT - 50b target |
| CGAGCTTGACGCATAGAGCGAGCGGCCCGCGCCTAAACATCCTGAAGCAC | Cas12a EMX1 - DT - 50b target - RC |
| GTGCTTCAGGATGTTTA**ATTAAAAATGAGCTTTTCTA**TGCGTCAAGCTCG | Cas12a CCR5 - DT - 50b target |

| | |
|---|---|
| CGAGCTTGACGCATAGAAAAGCTCATTTTTAATTAAACATCCTGAAGCAC | Cas12a CCR5 - DT - 50b target - RC |
| GTGCTTCAGGATGGCGC**CGGTTCGCGTGGATTAAAGG**TGCGTCAAGCTCG | Cas12a modified ps4 - DT1 - no PAM - 50b target |
| CGAGCTTGACGCACCTTTAATCCACGCGAACCGGCGCCATCCTGAAGCAC | Cas12a modified ps4 - DT1 - no PAM - 50b target - RC |
| AGCTTGTCTGCCATGGACATGCAGACTATACTGTTATTGTTGTACAGACCGAATTCCC | non-specific DNA target - no PAM |
| GGGAATTCGGTCTGTACAACAATAACAGTATAGTCTGCATGTCCATGGCAGACAAGCT | non-specific DNA target - no PAM - RC |
| GTGCTTCAGGATG<u>TTTA</u>**CGgTTCGCGTGGATTAAAGG**TGCGTCAAGCTCG | Cas12a modified ps4 - DT1 - 1MM - 50b target |
| CGAGCTTGACGCACCTTTAATCCACGCGAAACGTAAACATCCTGAAGCAC | Cas12a modified ps4 - DT1 - 1MM - 50b target - RC |
| GTGCTTCAGGATG<u>TTTA</u>**CGGTTCGCGTGGtTTAtAGG**TGCGTCAAGCTCG | Cas12a modified ps4 - DT1 - 2MM - 50b target |
| CGAGCTTGACGCACCTaTAAaCCACGCGAACCGTAAACATCCTGAAGCAC | Cas12a modified ps4 - DT1 - 2MM - 50b target - RC |
| GTGCTTCAGGATG<u>TTTA</u>**CGGTcCGCGTtGATTAgAGG**TGCGTCAAGCTCG | Cas12a modified ps4 - DT1 - 3MM - 50b target |
| CGAGCTTGACGCACCTcTAATCaACGCGgACCGTAAACATCCTGAAGCAC | Cas12a modified ps4 - DT1 - 3MM - 50b target - RC |
| GTGCTTCAGGATG<u>TTTA</u>**gGGTTCGtcTGGATTAgAGG**TGCGTCAAGCTCG | Cas12a modified ps4 - DT1 - 4MM - 50b target |
| CGAGCTTGACGCACCTcTAATCCAgaCGAACCcTAAACATCCTGAAGCAC | Cas12a modified ps4 - DT1 - 4MM - 50b target - RC |
| **Primers** | |
| AGAGTATAATGCTATTGTGGTTTTTGCGGATTTAAATTTTGGATTTAAAAGAGG | FnCas12a - DNase dead - E1006A - SDM primer 1 |

| | |
|---|---|
| CCTCTTTTAAATCCAAAATTTAAATCCGCAAAAACCACAATAGCATTATACTCT | FnCas12a - DNase dead - E1006A - SDM primer 2 |
| GCGGTTGTTGTCCTGGCGAACTTAAATTTTGG | AsCas12a - DNase dead - E993A - SDM primer 1 |
| CCAAAATTTAAGTTCGCCAGGACAACAACCGC | AsCas12a - DNase dead - E993A - SDM primer 2 |
| GTGATCGGAATTGCGCGTGGGGAGAG | LbCas12a - DNase dead - D832A - SDM primer 1 |
| CTCTCCCCACGCGCAATTCCGATCAC | LbCas12a - DNase dead - D832A - SDM primer 2 |
| CAATACCTAAAAAAATCACTGCGCCAGCTAAGAGGCAATAG | FnCas12a - pre-crRNA processing mutant - H843A - SDM primer 1 |
| CTATTGCCTCTTTAGCTGGCGCAGTGATTTTTTTAGGTATTG | FnCas12a - pre-crRNA processing mutant - H843A - SDM primer 2 |
| GAAGAACTGGTGGTGGCGCCCGCCAACAGCCCG | LbCas12a - pre-crRNA processing mutant - H759A - SDM primer 1 |
| CGGGCTGTTGGCGGGCGCCACCACCAGTTCTTC | LbCas12a - pre-crRNA processing mutant - H759A - SDM primer 2 |
| CACGTATGAAACGCATGGCCGCGCGTTTGGGAGAAAAG | AsCas12a - pre-crRNA processing mutant - H800A - SDM primer 1 |
| CTTTTCTCCCAAACGCGCGGCCATGCGTTTCATACGTG | AsCas12a - pre-crRNA processing mutant - H800A - SDM primer 2 |

| | |
|---|---|
| GTCAAGCTCGGACATCGTGATTGATAATGCGATGCACTGGCCGTCGTTTTACAACGTC | pUC19 plasmid library assembly - (between) M13 - vector amplification - common primer 1 |
| CCTGAAGCACCCTGTACGATTCGTACAGCAATGCGTCATAGCTGTTTCCTGTGTGAAATTG | pUC19 plasmid library assembly - (between) M13 - vector amplification - common primer 2 |
| TCGTCGGCAGCGTCAGATGTGTATAAGAGACAGGCATCGCATTATCAATCACGATGTC | for Nextera tagmentation - transposase adapter - forward |
| GTCTCGTGGGCTCGGAGATGTGTATAAGAGACAGGCATTGCTGTACGAATCGTACAGG | for Nextera tagmentation - transposase adapter - reverse |
